# Viral infection switches the balance between bacterial and eukaryotic recyclers of organic matter during algal blooms

**DOI:** 10.1101/2021.10.25.465659

**Authors:** Flora Vincent, Matti Gralka, Guy Schleyer, Daniella Schatz, Miguel Cabrera-Brufau, Constanze Kuhlisch, Andreas Sichert, Silvia Vidal-Melgosa, Kyle Mayers, Noa Barak-Gavish, J.Michel Flores, Marta Masdeu-Navarro, Jorun Karin Egge, Aud Larsen, Jan-Hendrik Heheman, Celia Marrasé, Rafel Simó, Otto X. Cordero, Assaf Vardi

**Author notes:** These authors contributed equally to this work (Flora Vincent, Matti Gralka).

## Abstract

Algal blooms are hotspots of marine primary production and play central roles in microbial ecology and global nutrient cycling. When blooms collapse, organic carbon is transferred to higher trophic levels, microbial respiration or sinking in proportions that depend on the dominant mortality agent. Viral infection can lead to bloom termination, but its impact on the fate of carbon remains an open question. Here, we characterized the consequences of viral infection on the microbiome composition and biogeochemical landscape of marine ecosystems by conducting a large-scale mesocosm experiment. Moniroting of seven induced coccolithophore blooms, which showed different degrees of viral infection, revealed that only high levels of viral infection caused significant shifts in the composition of free-living bacterial and eukaryotic assemblages. Intriguingly, viral infection favored the growth of eukaryotic heterotrophs (thraustochytrids) over bacteria as potential recyclers of organic matter. By combining modeling and quantification of active viral infection at a single-cell resolution, we estimate that viral infection can increase per-cell rates of extracellular carbon release by 2-4.5 fold. This happened via production of acidic polysaccharides and particulate inorganic carbon, two major contributors to carbon sinking into the deep ocean. These results reveal the impact of viral infection on the fate of carbon through microbial recyclers of organic matter in large-scale coccolithophore blooms.

## Introduction

Marine algae are responsible for half of Earth’s primary production and form the basis of the oceanic food chain^1^. Algal blooms^2^ are ephemeral events of phytoplankton proliferation that occur annually across the globe^3^ covering thousands of square kilometers. Upon demise, a small fraction of algal biomass is sequestered into the deep sea; the rest is transferred to higher trophic levels via predation, or recycled by heterotrophic bacteria and their predators, a process called the “microbial loop”^4,5^. It has long been hypothesized that the cause of bloom termination affects the surrounding microbiome and fate of carbon. Viral infection enhances lysis of host cells and release of dissolved organic matter (DOM), leading to bacterial growth at the expense of organic carbon sinking, in a process coined the “viral shunt”^6^. It has also been suggested that viral infection increases particle formation and thus biomass sinking. This accelerates the biologically driven sequestration of carbon into the deep sea in the so-called “viral shuttle” process^7,8^. However, we still lack quantitative assessment of how viruses alter microbial composition and influence the fate of carbon during algal blooms.

In order to provide a holistic and quantitative view of viral infection and its effect on carbon flow in the ocean, we investigated the dynamics of seven replicate mesocosm blooms of the cosmopolitan calcifying microalga *Emiliania huxleyi*. The blooms were induced by nutrient addition to a natural microbial community from the Norwegian fjord of Raunefjorden and the different enclosures spontaneously showed absent, moderate, or high levels of viral infection of the dominant alga. Combining daily monitoring of a variety of biological and biogeochemical parameters, we quantified the impact of viral infection on the associate eukaryotic and bacterial communities and on carbon cycling from the cellular to the biogeochemical level. Our results demonstrate how viruses impact microbial communities in coccolithophore blooms and their biogeochemical consequence on the fate of carbon.

### Succession of prominent community members during algal bloom dynamics

Our mesocosm experiment consisted of four uncovered enclosures (bag 1-bag 4) and three air-tight sealed enclosures to collect aerosols (bag 5-bag 7). The enclosures were immersed in a fjord near Bergen, Norway, filled with 11m^3^ of fjord water containing natural planktonic communities (**Fig. 1a**) and nutrients were added on series of consecutive days (**Fig. 1b**). During 24 days, we monitored phytoplankton and bacterial cell counts using flow cytometry, determined microbiome composition using amplicon sequencing (bacterial and eukaryotic), measured various biogeochemical parameters (see Methods), and determined infection states of single cells using single molecule fluorescent *in situ* hybridization^9^. These biological and chemical features were compared to the surrounding fjord waters.

**Figure 1:**
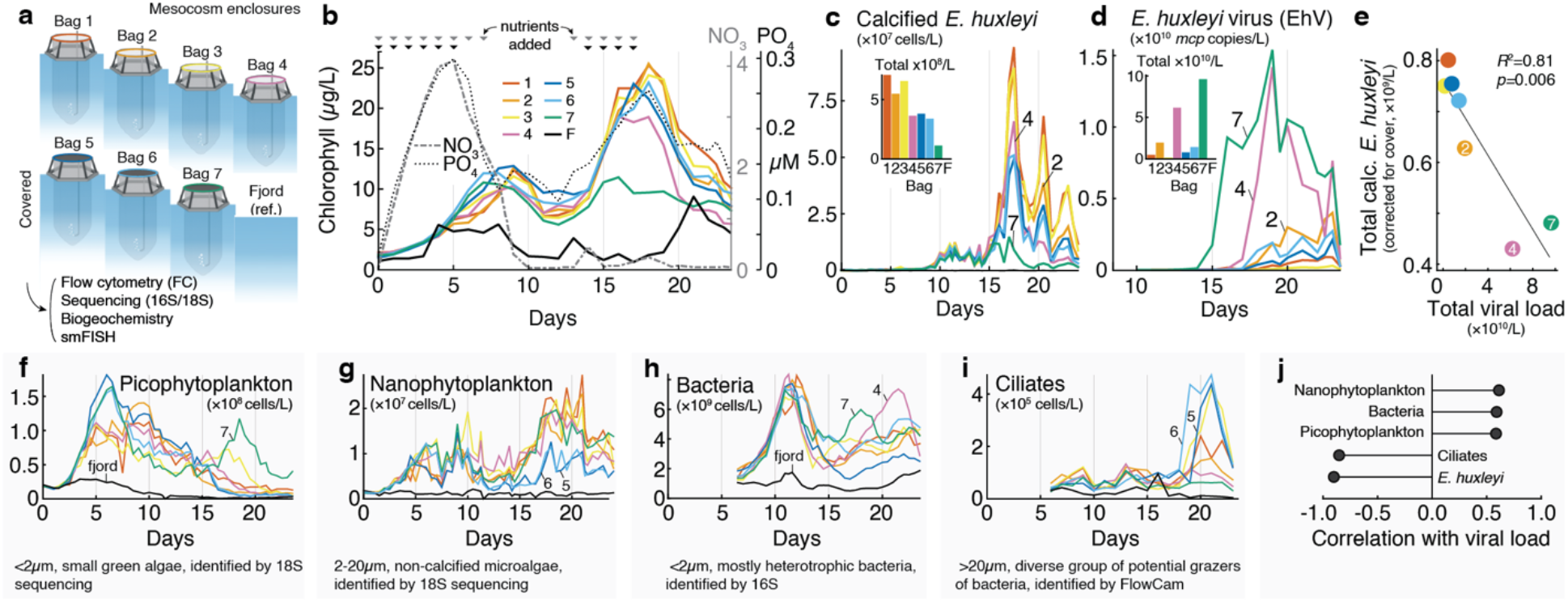
An overview of the mesocosm setup and dynamics of prominent community members. **a**, Schematic view of the seven mesocosm enclosures during the mesocosm experiment. The fjord water was used as the microbial inoculum seeding the enclosures and was sampled as a reference for microbial dynamics under natural conditions. **b**, Fluorometric chlorophyll measurements (left axis), where each color corresponds to a different bag. Nitrate and phosphate concentrations over time averaged across enclosures (right axes). Arrows on the top indicate nutrient addition. **c**, Calcified *E. huxleyi* abundance measured by flow cytometry, based on high side scatter and high chlorophyll signals. The small bar chart shows the integrated abundance of *E. huxleyi* over time (see Methods). **d**, Concentration of EhV based on qPCR of *mcp* (major capsid protein) gene in 2-20 μm pore filters. The small bar chart shows the integrated abundance of EhV over time. **e**, Scatter plot of total calcified *E. huxleyi* abundance (corrected for bag cover) as a function of total viral abundance, with a linear model fit. **f-i**, Absolute abundances of key players in the microbial succession, sorted by peak abundance time: **f**, picophytoplankton abundance measured by flow cytometry, based on low side scatter and low chlorophyll signals; **g**, non-calcifying *E. huxleyi* and other nanophytoplankton abundance measured by flow cytometry, based on low side scatter and high chlorophyll signals; **h**, absolute abundance of bacteria measured by flow cytometry after SYBR green staining; **i**, ciliate abundance measured by imaging flow microscopy and annotated using EcoTaxa. **j**, Correlation between EhV viral load and average planktonic abundances across bags.

In the initial phase (Day 0-10), the enclosures were nitrate and phosphate replete following nutrient addition; at later stages (Day 10-23), the enclosures were nitrate, but not phosphate, limited (**Fig. 1b).** Bulk chlorophyll measurements displayed two peaks in all enclosures, with a first smaller phytoplankton bloom (Day 0-10) followed by a second bloom reaching 25 μg/L of chlorophyll (Day 10-23). Calcified *E. huxleyi* cells, quantified by flow cytometry, dominated the second bloom (**Fig. 1c**). *E. huxleyi* is a cosmopolitan bloom-forming alga and one of the planet’s major calcite producers, causing the transport of large amounts of carbon into sediments^10^. In addition to slight differences in average *E. huxleyi* cell abundance between covered and uncovered enclosures, we also observed stark differences in *E. huxleyi* demise dynamics across bags (Day 18-24), with up to 90% lower algae abundances in bags 4 and 7 compared to the highest concentration within each day. Previous studies attribute the demise of natural blooms of *E. huxleyi* to virus-induced mortality caused by the *E. huxleyi*-specific giant coccolithovirus (EhV)^11–l3^. To assess the impact of EhV, we estimated its abundance in the 2-20 μm size fraction (to focus on infected *E. huxleyi* cells and viral particles associated with its biomass) by qPCR of the major capsid protein *(mcp)* gene (**Fig. 1d**). Viral abundance varied considerably between replicate mesocosms: bag 7 (covered) and bag 4 (uncovered) showed high concentrations of biomass-associated EhV with up to 1.54*10^10^ *mcp* copies/L and 1.42*10^10^ *mcp* copies/L, respectively, while bag 5 (covered) and bag 3 (uncovered) showed low to no detectable viral load. Virus-induced mortality had a direct impact on algal abundance: viral abundance explained 81% of the variance in *E. huxleyi* concentration across enclosures, suggesting that viral concentration controls the magnitude of an *E. huxleyi* bloom (**Fig. 1e**). For this estimate, we controlled for the different *E. huxleyi* abundances due to bag cover (bags 5-7) by adding the difference between covered and uncovered bag averages to the uncovered bags. Nonetheless, the fact that bloom demise was observed even with low or no viral infection suggested that other mortality agents may also dominate in *E. huxleyi* blooms. In enclosures with low viral load (bags 1, 3, 5, and 6), we observed up to a six-fold increase in ciliates that could potentially graze on *E. huxleyi*^14^ (measured by imaging flow microscopy, **Fig. 1i**).

The first phytoplankton bloom (Day 0-10) which we termed the mixed bloom, preceding the *E. huxleyi* bloom, was dominated by the pico-phytoplankton *Bathycoccus* and *Micromonas,* representing over 40% of the community in the 0.2-2 μm size-fraction (**Supplementary Fig. 1**). This bloom reached 1.81*10^8^ cell/L in bag 5 (**Fig. 1f**). Nano-phytoplankton (**Fig. 1g**) were also important players in this mixed bloom and sequencing of the 2-20 μm size fraction 18S rDNA revealed that dinoflagellates (Group-I Clade-I) were especially abundant (see further information below).

Phytoplankton cells fix inorganic carbon into organic biomass, and secrete part of it in the form of metabolites that heterotrophic bacteria can use for growth^15–18^. Interestingly, the dissolved organic carbon (DOC) concentration increased only moderately after each of the blooms (**Supplementary Fig. 2**). This could be explained by a fast-bacterial assimilation as we observed a more than tenfold exponential increase in bacterial abundance between days 5-13 (**Fig. 1h**), doubling every 24-36 hours. By contrast, bacteria were less abundant during the *E. huxleyi* bloom and demise compared to the mixed bloom, showing an average of less than twofold increase after day 20 (**Fig. 1h**). In the two most infected bags, bag 4 and bag 7, the increase in bacterial abundance was 2-3-fold during the demise phase. Overall, total viral load in the different enclosures was significantly negatively correlated with the abundance of host (*E. huxleyi*) and grazer (ciliates) concentrations but not with pico-nano-phytoplankton or bacteria abundances (**Fig. 1j**). The negative correlation between grazing and viral lysis was confirmed via grazing dilution assays across the whole mesocosm (**Supplementary Fig. 3**), suggesting that the two top-down mortality agents compete during algal blooms.

### Effects of viral infection on the composition of microbial assemblages

To understand how viral infection can alter the composition of planktonic communities, we conducted microbiome profiling of all the mesocosm enclosures. We opted for a detailed time series based on samples collected daily of both the bacterial (0.2-2 μm size fraction, 16S amplicons) and nanoeukaryotic (2-20 μm size fraction, 18S amplicons) communities. Throughout the two blooms, we observed a repeatable pattern of eukaryotic and bacterial taxa successions (**Fig. 2a**). The relative abundance of *E. huxleyi* defines three major phases: the mixed bloom, (days 0-8), the exponential growth phase of *E. huxleyi* (days 8-17), and its demise (days 18-23). Nanoeukaryotes, clustered according to the relative abundance patterns at the genus level, showed a rapid succession of boom-and-bust cycles, each about 5-10 days long (**Fig. 2b, Supplementary Fig. 4,5**). Unique clusters of nanoeukaryote species bloom upon *E. huxleyi* growth (Cluster 5) and demise (Cluster 6) thus defining a bloom associated protist microbiome. By comparison, bacterial succession was much less dynamic: the composition at the order level was relatively stable (**Fig. 2c**). The mixed bloom was associated with bacterial groups known to be involved in algal biomass remineralization, such as Flavobacteriales^19,20^ and Rhodobacterales. We also observed a slow, more than ten-fold increase in relative abundance of SAR11, usually found in oligotrophic environments^21^, throughout the *E. huxleyi* bloom. This facilitation of SAR11 growth by *E. huxleyi* is in line with previous observations of their co-occurrence^22^ and could be mediated by the organosulfur compound dimethylsulfoniopropionate (DMSP), which *E. huxleyi* produces and excretes^23,24^ and that SAR11 can utilize as a reduced source of sulfur^25^. In contrast to the relative stability of the bacterial composition at the order level, there were clear successions at the genus level within the two dominant bacterial orders, *Flavobacteriales* and *Rhodobacteriales* (**Fig. 2d, Supplementary Fig. 6**). The *E. huxleyi* bloom and demise coincided with the relative increase of two genera: *Tenacibaculum*, a potential fish parasite frequently associated with algal blooms^20^, and *Sulfitobacter*, a genus containing DMSP degrading species that are pathogenic to *E. huxleyi* cells^24^. Gammaproteobacteria such as *Vibrionales*, *Pseudomonadales*, or *Alteromonadales*, often reported as dominant members in bloom-associated communities^26^, were absent in the planktonic populations, but may thrive in the particle-associated niche (in the >20 μm size fraction).

**Figure 2:**
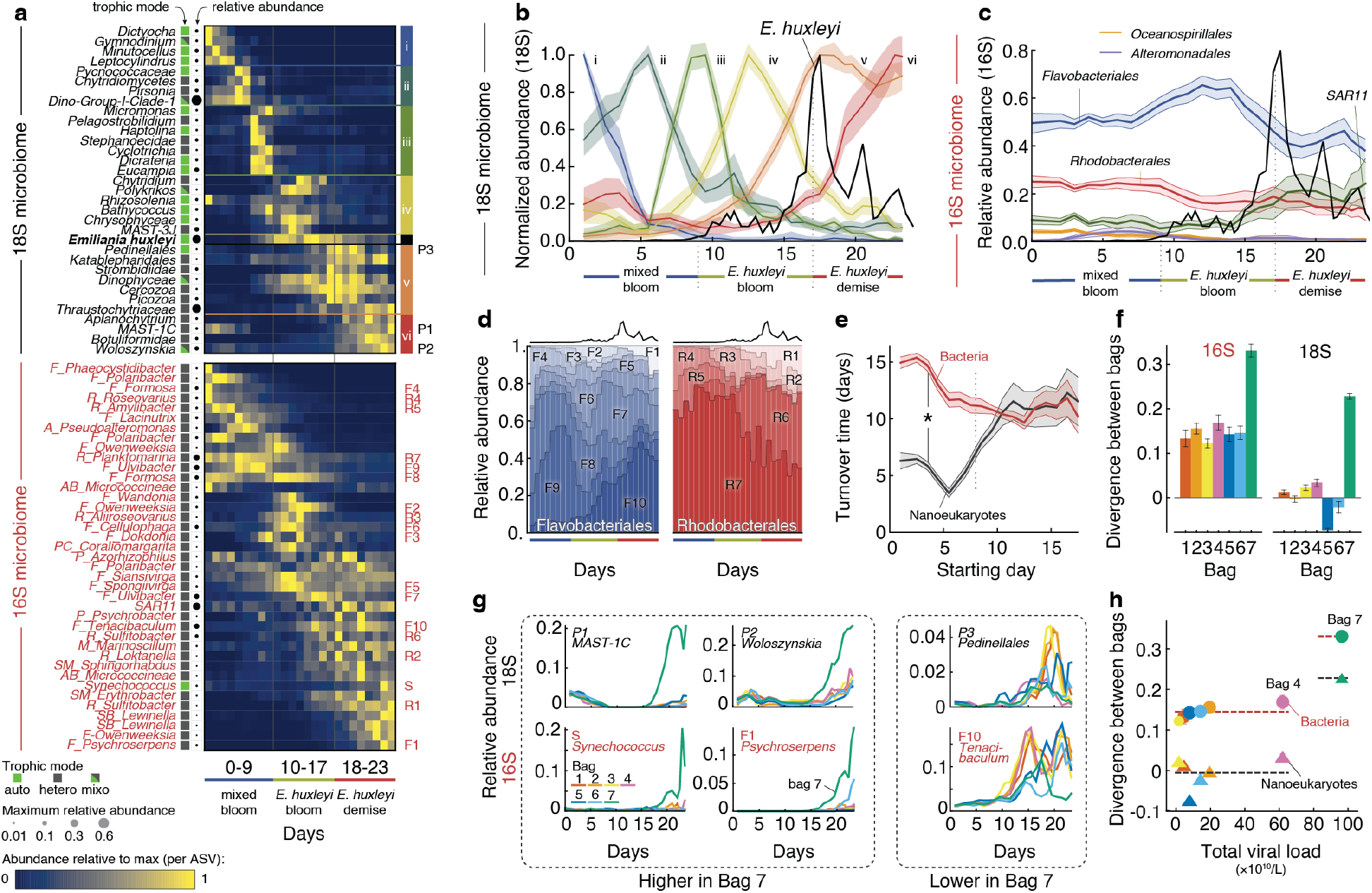
Microbial succession during the growth and demise of algal blooms with different viral loads. **a**, Bacterial and eukaryotic microbial succession throughout the experiment duration, averaged across enclosures. Each row is an amplicon sequencing variant (ASV) with bacteria in red and eukaryotes in black. The trophic modes of each ASV are detailed in the box color with autotrophs in green, heterotrophs in grey, and mixotrophs in green/grey. Days are shown in columns. 18S species are grouped by clusters of different colors, indicated on the right of the heatmap. 16S abbreviations: F: Flavobacterales; R: Rhodobacterales; A: Alteromonadales AB: Actinobacteridae; M: Methylophylales; P: Pseudomonadales; PC: Puniceicoccales; SM: Sphingomonadales; SB: Sphingobacteriales. **b**, Succession of 18S-based ASVs in the 2-20μm fraction, clustered by similarity of their relative abundance dynamics averaged across bags. The shaded area represents the standard deviation within each cluster. The absolute abundance of *E. huxleyi* enumerated with flow cytometry is overlaid as a guide (black line, not to scale). Each cluster is normalized to its own maximum abundance and their species composition is detailed in panel **a**. **c**, Relative abundance of major bacterial orders throughout the bloom, averaged for all enclosures using 16S amplicon sequencing of the 0.2-2 μm fraction. The absolute abundance of *E. huxleyi* enumerated with flow cytometry is overlaid as a guide (black line, not to scale). **d**, Relative abundance of different Flavobacteria and Rhodobacteria genera within each order averaged across all enclosures. The dark line on the top represents *E. huxleyi* abundance trends as a guide. **e**, Rate at which bacteria and nanoeukaryotic community similarities change over time. Nanoeukaryotic communities initially turnover much faster than the bacterial ones (until day 8, p<0.001 by Kolmogorov-Smirnov test). **f**, Compositional divergence within one bag, compared across several days, for 16S and 18S. The divergence of a bag is defined as the change in pairwise Bray-Curtis distance between the focal bag and all other bags from the start of the *E. huxleyi* bloom to its demise. **g**, Eukaryotic and bacterial ASVs that are overrepresented or underrepresented in bag 7. **h**, Correlation, per bag and per 16S or 18S ASV, between total viral load and percentage dissimilarity in microbial composition from one day to another.

To further compare bacterial and eukaryotic dynamics, we computed their turnover time as defined by the exponential rate at which the Bray-Curtis similarity declined over time (see Methods). Given their small size and known fast growth rates, we expected heterotrophic bacteria to respond much faster to our nutrient additions (N and P) than eukaryotes. To our surprise, eukaryotes were the first responders to nutrient addition, and their assemblage turned over much faster (every five days initially) than bacteria which only showed significant growth towards the end of the first bloom (turnover every 10 days) (**Fig. 2e**). The sequence of response to the nutrient addition can be explained by the direction of nutrient flow in phytoplankton blooms when nutrients increase: eukaryotes, especially phytoplankton, were likely nitrogen and/or phosphorous-limited at the start of the experiment, whereas bacteria appeared to be carbon-limited and required organic carbon released upon demise of the first mixed bloom in order to grow.

Despite strong compositional similarities amongst the seven enclosures, the bacterial and nanoeukaryotic assemblages gradually diverged between enclosures after the mixed bloom. During the *E. huxleyi* bloom demise, bag 7 (the most virally infected) diverged in microbiome composition from the other enclosures (**Fig. 2f, Supplementary Fig. 7**). Eukaryotes such as MAST-1C (a heterotrophic flagellate), *Woloszynskia* (a mixotrophic dinoflagellate), as well as the cyanobacterium *Synechococcus* and the bacterium *Psychroserpens* (family *Flavobacteriaeceae*) were overrepresented in bag 7 (**Fig. 2g, Supplementary Fig. 8,9**). The growth of *Synechococcus* during high viral infection suggests that the resulting flux of DOM benefit not only heterotrophic but also autotrophic bacterial growth^27,28^. Recent ecosystem modeling suggests this may be due to efficient recycling of growth-limiting nutrients in the photic zone during viral infection^29^. The eukaryotes *Pedinellale* (autotroph) and the bacterium *Tenacibaculum* (family *Flavobacteriaeceae*) grew less in bag 7 that in the rest of the enclosures. In contrast to observation in the highly infected bag 7, the moderately infected bag 4 showed a 16S and 18S-based composition that did not differ significantly from the less infected enclosures (**Fig 2h**). These findings suggest that substantial change in microbial assemblages during viral induced *E. huxleyi* demise is conditional on high viral infection levels.

### Viral infection impacts the composition of organic matter recyclers

During *E. huxleyi* demise, a large flux of organic carbon derived from lysed phytoplankton biomass became available for bacterial recycling, with estimates of about 270 μg C/L/day from *E. huxleyi* alone (see Methods). Yet bacterial growth was moderate. This could be explained by several factors, including enrichment of particle-attached (e.g., biofilm-like) bacterial growth that did not influence the free-living abundances, removal of bacteria by aggregation and sinking, or increased bacterial cell death by phages or bacterivores^30^. However, the abundance of typical bacterivores like dinoflagellates remained low and ciliate abundance only increased late into the demise of the *E. huxleyi* bloom (day 20-23) (**Fig. 1i**). The low number of predators, combined with the observation that dissolved organic carbon concentration stabilized during bloom demise, led us to hypothesize that bacteria competed for nutrients with another group of heterotrophs.

To identify other heterotrophs, we re-examined the eukaryotic microbiome in search for organic matter recyclers. Functional annotation of the nanoeukaryotes (see Methods) revealed that while eukaryotic assemblages were composed of autotrophs and mixotrophs during the first mixed bloom, heterotrophs, and specifically osmotrophs, became highly abundant through the *E. huxleyi* bloom and demise (**Fig. 3a**). These heterotrophs were dominated by thraustochytrids (*Thraustochytriaceae* and *Aplanochytrium* in **Fig. 2a**), part of a diverse lineage of eukaryotic osmotrophs^31^, which contributed over 50% of all 18S rDNA reads in the 2-20 μm size fraction during bloom demise, across all bags. Thraustochytrids are known to possess an arsenal of extracellular digestive enzymes, making them important decomposers of organic matter in coastal sediments^32^ and deep-sea particles^33^. With their large intracellular lipid reserves, they also serve as an important food source for higher trophic levels^34^. However, the importance of thraustochytrids in microbial food webs has yet to be explored. During algal blooms, they could potentially play a significant role as decomposers^35^, bacterivores, or even parasites^36^. Some members of the group are also known to produce ectoplasmic nets, through which they can extract intracellular nutrients of preyed cells^37,38^ such as senescent diatoms^39,40^.

**Figure 3:**
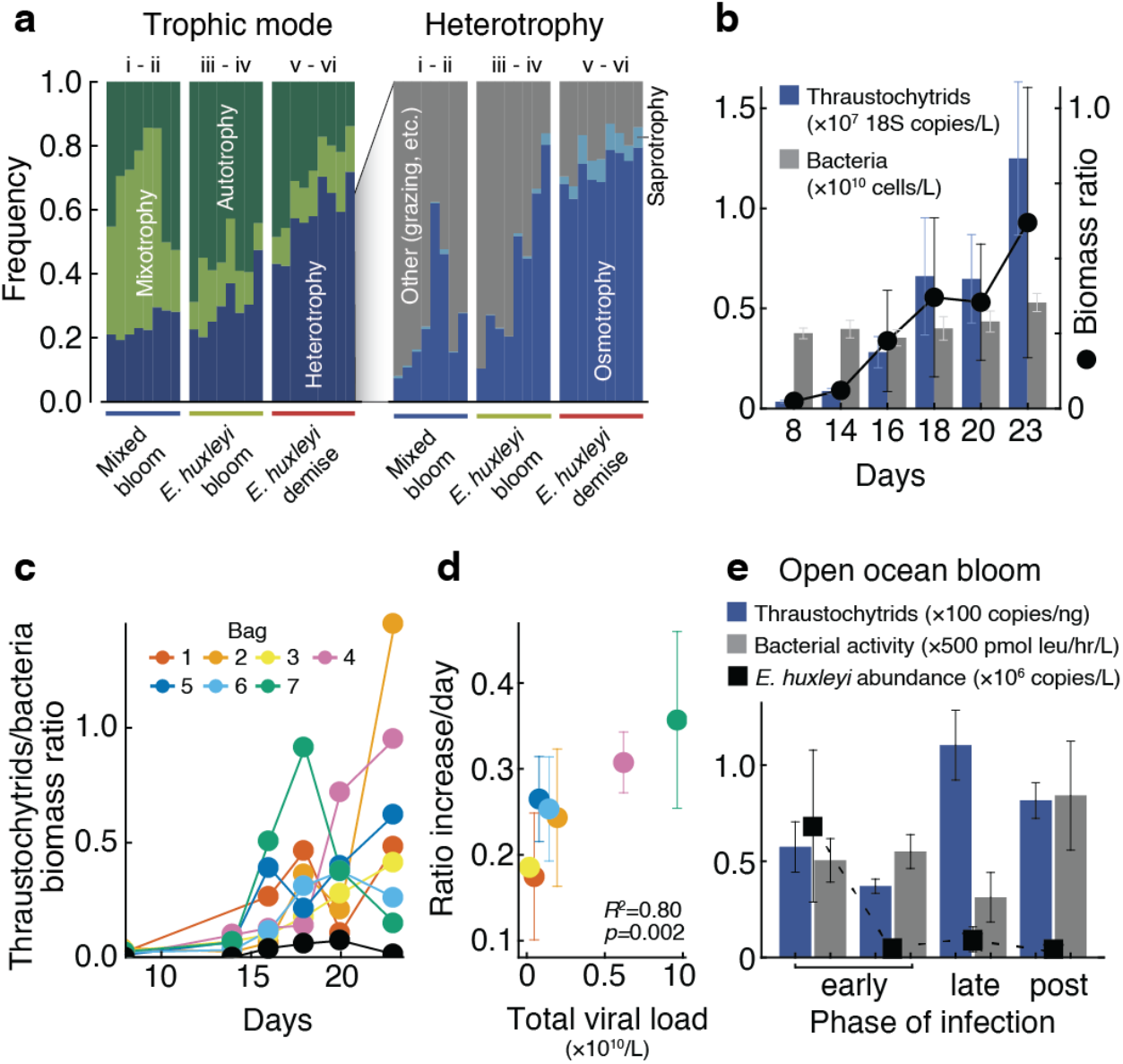
Viral induced bloom demise changes the composition of organic matter recyclers. **a**, Analysis of predicted eukaryotic traits plotted per phase of the *E. huxleyi* bloom. Relative abundance of autotrophy, mixotrophy, and heterotrophy, defined through literature search (left). Trophic modes within heterotrophy, as defined by^45^ (right). **b**, Thraustochytrid to bacteria ratios throughout the experiment duration. The blue bars represent thraustochytrid concentrations obtained by digital droplet PCR and the grey bars represent bacterial concentrations measured by flow cytometry, averaged across all bags. Circles represent ratios of thraustochytrids to bacterial biomass in the averaged mesocosm enclosures, using a conservative conversion factor of thraustochytrid copies/L to cell/L (see Methods and **Supplementary Table 1**). **c**, Converting thraustochytrid and bacteria abundances to biomass to obtain biomass ratios shows increasing ratio during *E. huxleyi* bloom (days 10-17) and demise (days 17-24) in all bags. **d**, Exponential rate of change of the biomass ratio of thraustochytrids to bacteria plotted as a function of total viral load, per bag (see Methods). **e**, Concentration of thraustochytrids measured by ddPCR, *E. huxleyi* cells measured by qPCR, and bacterial production using leucine incorporation^44^, in three phases of an open ocean *E. huxleyi* bloom infection.

In order to quantify the absolute abundance of thraustochytrids, we performed digital droplet PCR (ddPCR) targeting thraustochytrid 18S rDNA across all mesocosm enclosures. While undetected during the mixed bloom, thraustochytrids total biomass, estimated based on values of 64 18S rDNA copies/cell (see Methods) and 1.65×10^−10^ g of carbon/cell^41^ increased steadily after day 16 and was comparable to that of bacteria during the *E. huxleyi* bloom demise (**Fig. 3b, Supplementary Table 1**). Though we cannot elucidate the mechanism of competition between thraustochytrids and heterotrophic bacteria, possibilities include direct inhibition of bacterial growth by antimicrobial lipids^42^, niche separation in the degradation of different components of the organic matter, or efficient capture of organic matter by ectoplasmic nets directly from senescent *E. huxleyi* cells. Thraustochytrids are most likely not intracellular parasites of *E. huxleyi*, since we did not detect any 18S amplicon reads of this group within *E. huxleyi* cells that were sorted and sequenced.

While the *E. huxleyi* demise reproducibly triggered the growth of eukaryotic degraders in all bags, thraustochytrids growth was further enhanced in bags with strong viral infection of *E. huxleyi* (**Fig. 3c**). Specifically, total viral load was well correlated with the exponential rate at which the ratio of thraustochytrid to bacterial biomass increased (**Fig. 3d**). We further examined the ecological importance of this phenomena by using samples collected from an open-ocean *E. huxleyi* bloom in the North Atlantic^43^ where different phases of viral infection were observed. We quantified the absolute abundances of thraustochytrid using ddPCR and compared it with bacterial production rates measured with leucine incorporation^44^. We detected higher thraustochytrid abundance and lower bacterial production during late viral infection phase relative to the early phase of bloom infection (**Fig. 3e**), suggesting that thraustochytrids are major benefiters from viral induced *E. huxleyi* bloom demise. Sequencing of larger 18S rDNA fragments from the mesocosm and open ocean samples revealed a single dominant species across these ecosystems, whose closest relative is an uncultivated clone (94% identity), potentially indicating that this thraustochytrid species specializes on exudates from *E. huxleyi* demise and has not been reported before (**Supplementary Fig. 10**).

### Viral infection enhances population-level and per-cell rates of carbon release

During *E. huxleyi* blooms, which can cover over 100,000 square kilometers in the ocean^46^, cell concentrations can account for 75% or more of the total number of photosynthetic plankton in the area^46^. The algal biomass and coccoliths that form the *E. huxleyi’s* calcified shell have profound impact on the carbon cycle and global CaCO_3_ export flux^47^. We therefore investigated the biogeochemical consequences of viral infection of *E. huxleyi* blooms by quantifying different components of the carbon cycle, focusing on organic carbon in the form of transparent exopolymer particles (TEP) and particulate inorganic carbon (PIC).

TEP are made of acidic polysaccharides that form due to abiotic coagulation of dissolved carbohydrates secreted by phytoplankton and are an important component of the marine particulate organic carbon. TEP represent a potential source of food for bacteria or other heterotrophs^48,49^, although recent work suggests that certain polysaccharides within TEP can be recalcitrant for microbes, challenging its degradation^50^. TEP are an essential vector for carbon export by triggering aggregation and sinking but their chemical composition has only recently been elucidated. To better identify the polysaccharides in TEP during *E. huxleyi* blooms, we used carbohydrate microarray analysis on the particulate fraction^51^. Out of the alginate and sulfated fucans epitopes, the ones recognized by the monoclonal antibodies BAM6^52^ and BAM1^53^ respectively, accumulated during the *E. huxleyi* bloom and are thus likely *E. huxleyi*-related (**Fig. 4a**). BAM6 signal decreased during the demise phase, suggesting potential degradation of its recognized epitope by the demise associated microbiome. In contrast, the accumulation of the epitope detected with BAM1 suggests that this sulfated fucan did not serve as a substrate for thraustochytrids or bacteria^54^, but may be part of TEP and thus relevant for carbon export via sinking particles^50^.

**Figure 4:**
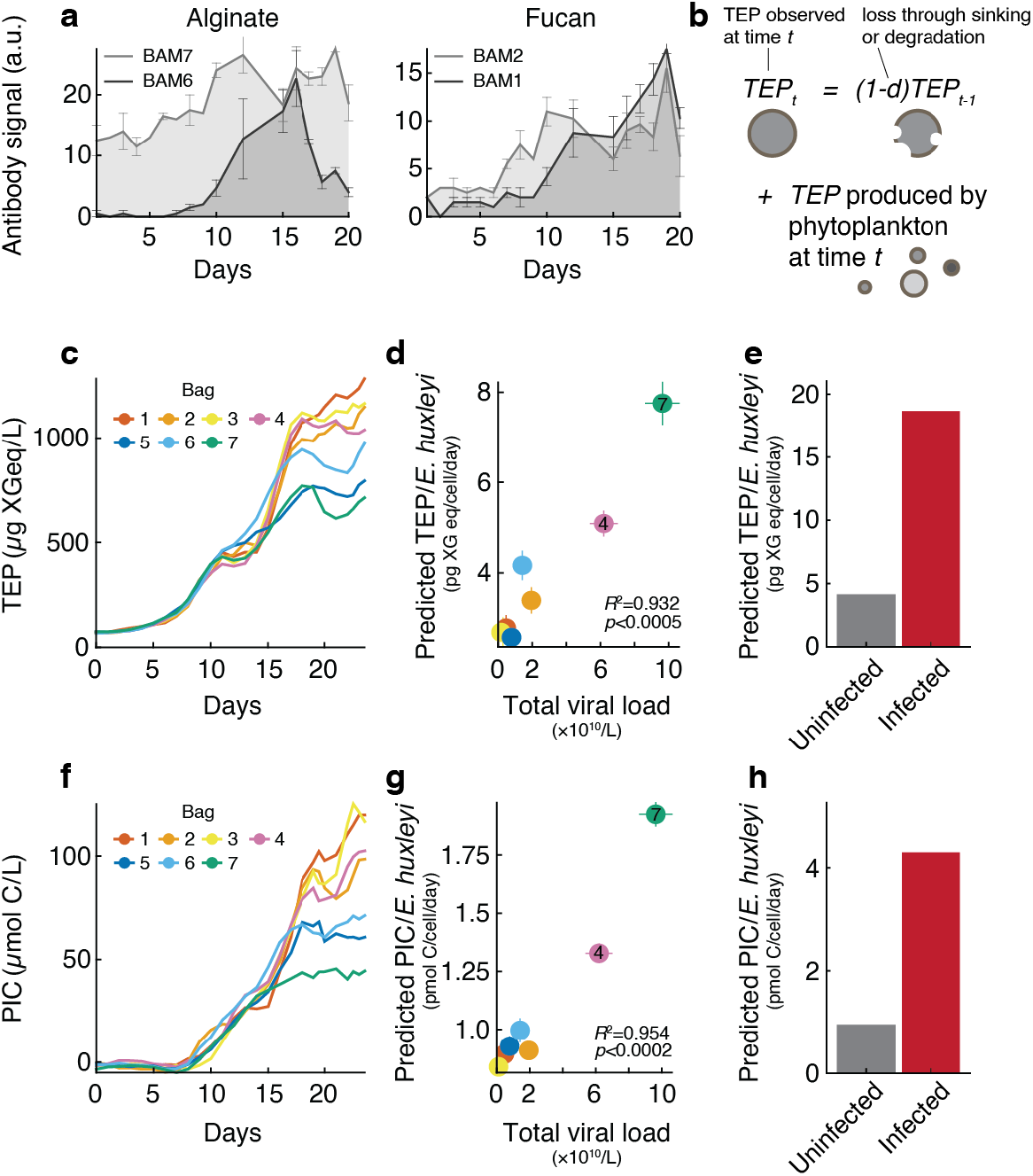
Viral infection promotes release of PIC and TEP production from a coccolithophore bloom. **a**, Alginate and fucan abundance in particulate organic matter (POM) over time, based on mixed water from bags 1-4, measured by carbohydrate microarray analysis. BAM1, 2, 6 and 7 correspond to glycan-specific monoclonal antibodies, used to measure the relative abundance of their recognized polysaccharide epitopes in POM water extracts. **b**, Scheme of TEP modeling, as a function of phytoplankton concentrations and degradation rate which enables prediction of the *E. huxleyi* contribution to the TEP pool. **c**, TEP concentration measured by Alcian blue staining over time, per bag. **d**, Predicted TEP/cell as a function of total viral load for each bag, for *E. huxleyi* cells. **e**, Predicted TEP/cell secretion in infected versus non-infected *E. huxleyi* cells using intracellular measurements of actively infected single cells. **f**, PIC production during algal bloom succession. **g**, Predicted PIC/cell as a function of total viral load for each bag, for *E. huxleyi* cells. **h**, Predicted PIC/cell production in infected versus non-infected *E. huxleyi* cells using intracellular measurements of actively infected single cells.

To decipher the effects of viral infection on TEP production, we modeled TEP concentration as a function of its producers’ abundances (*E. huxleyi*, non-calcified nanophytoplankton and picophytoplankton) and a TEP loss rate through sinking or degradation (**Fig. 4b**) that we fitted to *in situ* TEP measurements (**Fig. 4c**) (see Methods). The model described the TEP abundance well, achieving an average *R^2^* of 98.8%. Using the model, we estimated that the amount of TEP produced per *E. huxleyi* cell per day was 60-75% of the total TEP pool at the onset of bloom demise (**Supplementary Fig. 11**). There was a strong dependence of estimated TEP per cell on viral infection: TEP production per *E. huxleyi* cell was more than twice as high in the infected bag 7 than in non-infected bags. Across all bags, there was a strong correlation to total viral load (*R*^2^ = 0.932,*p* < 0.0005, **Fig. 4d**), consistent with previous results suggesting higher export during viral-associated *E. huxleyi* blooms in open ocean and mesocosm experiments^43,55^. This suggests that, at the population level, *E. huxleyi* cells secreted twice the amount of carbon in presence of high viral load. To validate this correlation, we applied the same model for particulate organic carbon (POC) production. The model gave an excellent fit (*R^2^* >0.98 across all bags) but the estimate for the amount of organic carbon per *E. huxleyi* cell (4-6 pg C/cell, in line with other estimates^56^ (**Supplementary Fig. 12**)) was uncorrelated with the total viral load (*p*>0.05).

Since viral infection remodels the algal host metabolism^57,58^, we hypothesized that infected and non-infected cells in the same bloom may differ in their actual TEP production, and sought to quantify this process as opposed to simply averaging TEP over the entire bulk population. To differentiate infected from non-infected cells, we probed viral mRNA in single *E. huxleyi* cells^9^ and obtained a time-course of the fraction of actively infected cells in two different enclosures (**Supplementary Fig. 13**). At most 10% and 25% of all *E. huxleyi* cells were infected in bags 2 and 4 respectively, reflecting the heterogeneity of cell fates within each bloom succession and demise. By assuming that non-infected cells produced the same amount of TEP regardless of the bag’s viral load, we estimated that an infected *E. huxleyi* cell produced ~19 pg xanthan gum (XG) equivalent/day (see Methods), or 4.5 times more TEP than its non-infected bystander cell (**Fig. 4e**). Notably, viral infection did not increase secretion of proteinaceous material: the measurement and modeling of protein-rich particles (Coomassie Stained Particles) (**Supplementary Fig. 14**) showed no correlation with viral load, indicating that the cellular response to infection is specific to certain metabolic products.

Particulate inorganic carbon (PIC) in the form of calcium carbonate is the basis of one of the main processes making up the marine carbon cycle, the carbonate pump, by which inorganic carbon is exported along with organic matter to the deep ocean. A major part of PIC in the ocean is comprised of coccolithophore shells, particularly *E. huxleyi’s* coccoliths^59^. PIC accumulated over time in our study (**Fig. 4f**). We fitted the PIC curves to a model accounting for *E. huxleyi* coccolith production, a degradation rate, and a term allowing for shedding and re-calcification (see Methods). Like TEP, predicted PIC per *E. huxleyi* cell at the population level was significantly correlated to total viral load from about 1 to 2 pmol PIC/cell/day (*R*^2^ = 0.954,*p*<0.0002, **Fig. 4g**) which is consistent with lab-based measurements^60^. Using the measured fraction of active single-cell infection, we estimated that infected single cells produced 4.5 times more PIC per cell than their non-infected bystander cells (**Fig. 4h**). Overall these data suggest that active viral infection can have remarkable consequences on exportable carbon (TEP and PIC) release both on the population-level (2-fold increase) and per infected cell (4.5-fold increase).

### Conclusion

Here we provide in depth characterization of the microbial and biogeochemical dynamics of two successive algal blooms in replicate mesocosm enclosures, which provides a unique experimental platform to quantify the impact of viral infection at the ecosystem level. Starting from the same microbial inoculum, our mesocosm enclosures underwent ordered microbial successions that culminated in massive blooms of the coccolithophore *E. huxleyi*. Viral infection of *E. huxleyi* took drastically different courses in the enclosures, with little to high levels of viral infection leading to a bloom demise. Our study made three critical observations regarding the microbial ecology and the biogeochemical effects of algal blooms and their viral infection (**Fig. 5**), generating novel hypotheses for future lab-based mechanistic studies.

**Figure 5:**
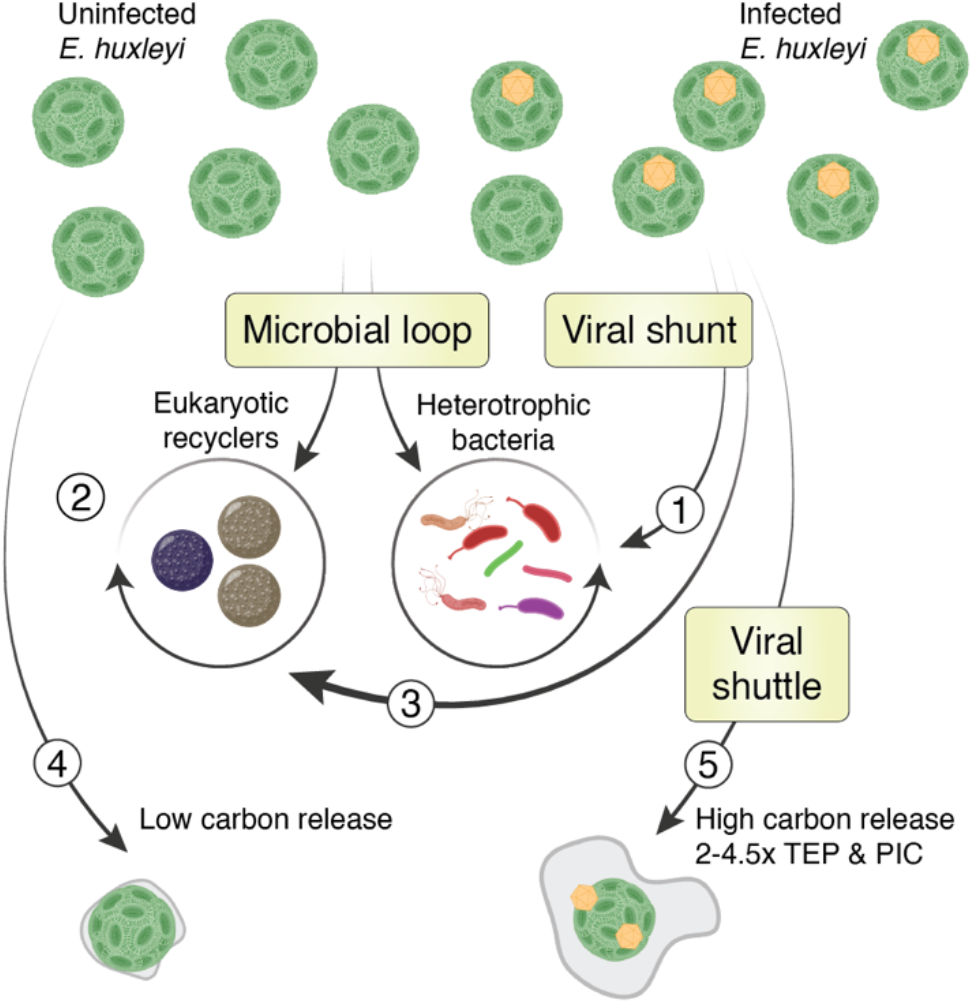
Consequences of viral infection on microbial community composition and carbon cycling. Arrows represent the direction of carbon flow. (**1**) The bacterial and eukaryotic microbiomes are remodeled in response to viral infection only when level of infection is high. (**2**) Thraustochytrid rival bacteria as significant recyclers of organic matter during *E. huxleyi* demise. (**3**) Thraustochytrids benefit from viral infection of *E. huxleyi*. (**4**) When the demise is not virus-associated, *E. huxleyi* populations release a small amount of organic and inorganic carbon. (**5**) Viral infection increases *E. huxleyi* population carbon release between 2-4.5 fold under the form of TEP and PIC as compared to (**4**).

First, we showed that viral-induced changes in the microbiome, as in bag 7, are only observed when there is a high level of viral infection (**Fig. 5** (**1**)). Given that only one in seven enclosures experienced such high levels of viral production, its occurrence in natural ecosystems may be rare but can profoundly impact microbial diversity and community composition. It is also possible that the microbiome response takes longer than the duration of our experiment or may be localized to particle-attached communities. Indeed, as viral infection enhanced TEP production, this could promote the formation of marine snow and a specific particle-associated microbiome.

Second, we estimated that the biomass of eukaryotic osmotrophs can be comparable to that of heterotrophic bacteria during *E. huxleyi* blooms and demise (**Fig. 5** (**2**)) and that viral infection may enhance their growth (**Fig. 5** (**3**)), including in open ocean blooms. This abundance of large, underappreciated eukaryotic osmotrophs, may shape the carbon flux through the marine trophic network. Since they are larger than the average bacterium, eukaryotic osmotrophs can escape grazing by many micrograzers and shorten the carbon transfer to larger predators as zooplankton. Thus, the competition between prokaryotic and eukaryotic degraders can reshape higher trophic levels. More generally, our findings highlight that a complete understanding of the carbon flux in phytoplankton blooms requires deeper understanding of both the associated prokaryotic and eukaryotic microbial communities and the interactions between them^61^. Currently, only a few studies explore the competition between thraustochytrids and bacteria for DOM^62^. More work is needed to fully establish the role of eukaryotic osmotrophs in the microbial loop, especially in the context of viral infections and the associated metabolome^63^.

Third, by relating ecosystem changes and biogeochemical processes to the varying degrees of active viral infection and lysis, we have shown that even mild viral infection can significantly affect the production and release of extracellular carbon, both organic and inorganic. In particular, our experimental setup enabled the parameterization of TEP and PIC production in the absence (**Fig. 5** (**4**)) or presence (**Fig. 5** (**5**)) of viral infection^43,55,64^. We estimated that *E. huxleyi* carbon secretion in the presence of viruses increases between 2 to 4.5-fold per cell, either through a population level response, or a specific metabolic remodeling of infected cells. Taken together, the increase in TEP and PIC production per cell could lead to elevated vertical carbon transport through aggregation and increase of the cellular ballast. Overproduction of TEP by infected cells increases the formation of sinking aggregates, and may protect non-infected cells by trapping newly produced virions in sticky particles or can mask receptors needed for viral entry^9^. Alternatively, TEP could be involved in the transport of virions to neighboring cells, in analogy to the human T-cells leukemia virus which encases itself in a host-derived carbohydrate-rich adhesive extracellular cocoon that enables its efficient and protected transfer between cells^65,66^. The increased production of PIC per cell is surprising since viral infection is thought to promote decalcification^67^. Nevertheless, higher turnover of coccolith shedding and recalcification, or thicker coccoliths could be potential defense mechanisms, enabling lower viral adsorption and efficient removal of attached viral particles.

Taken together, our results provide a strong evidence that viral infection does not only play an important ecological role as a principal cause of phytoplankton mortality, but also has profound consequences for the fate of carbon, both by diverting carbon from bacteria towards larger eukaryotes and by potentially enhancing vertical export (**Fig. 5**). This refined assessment of viral impacts on the fate of carbon in the ocean helps bridge the scales between dynamic processes at the single cell, population, and biogeochemical levels, and will thus enable us to anticipate better the consequences of a changing ocean on fundamental ecosystem processes, services and feedbacks.

## Acknowledgments

We thank all team members of the AQUACOSM VIMS-Ehux project for setting up and conducting the mesocosm experiment. We further thank the team members and crew of the NA-VICE cruise for assistance at sea, as well as the Marine Facilities and Operations at the Woods Hole Oceanographic Institution for logistical support. We thank Miri Shnayder for help on ddPCR, Tina Trautmann for help on microarrays and Jackie Collier for discussion on Thraustochytrids.

## Funding

A.V. is The Bronfman Professorial Chair of Plant Science. This research was supported by the European Research Council CoG (VIROCELLSPHERE grant no. 681715), the research grant from the Estate of Bernard Berkowitz and the Simons foundation grant (no. 735079) “Untangling the infection outcome of host-virus dynamics in algal blooms in the ocean” awarded to A.V. and the Dean of Faculty Fellowship of the Weizmann Institute of Science and Israeli Academy of Science and Humanities awarded to F.V. The mesocosm experiment VIMS-Ehux was supported by EU Horizon2020-INFRAIA project AQUACOSM (grant no. 731065). The NA-VICE cruise was supported by the NSF (grant no. OCE-1059884). J.-H.H. was supported by the Max Planck Society and by the Deutsche Forschungsgemeinschaft (DFG) Emmy Noether grant HE 7217/1-1, and through the Cluster of Excellence “The Ocean Floor — Earth’s Uncharted Interface” project 390741603. O.X.C. was supported by the Simons Collaboration: Principles of Microbial Ecosystems (PriME) award no. 542395. M.G. was supported by Simons Foundation Postdoctoral Fellowship Award 599207. The ICM-CSIC group acknowledges funding from Spanish Ministry of Science and Innovation (MCIN/AEI, doi: 10.13039/501100011033) through the BIOGAPS grant (CTM2016–81008–R) and the “Severo Ochoa Centre of Excellence” accreditation (CEX2019-000298-S), and from the European Research Council under the EU’s Horizon 2020 research and innovation programme through the SUMMIT grant (ERC-2018-AdG#834162).

## Author contributions

F.V., M.G., and A.V. conceptualized this study and wrote the manuscript, with input from J.-H.H., R.S. and O.X.C. F.V. and M.G. acquired flow cytometry data, performed amplicon sequencing, ddPCR analysis, smFISH, designed and wrote all scripts for data analysis. G.S., D.S. performed the qPCR analysis. M.C., M.MN., and C.M. collected biogeochemical data. C.K. and K.M. collected Flowcam data. A.S., and S.V.-M. performed epitope analysis. K.M. performed grazing assays. N.B.G. and M.F., performed biomass filtration. J.K.E. and A.L. assisted during the mesocosm experiment. All authors reviewed and edited the manuscript.

## Competing interests

The authors declare that they have no competing interests.

## Data and materials availability

All data needed to evaluate the conclusions in the paper are present in the paper and clearly indicated in the Supplementary Materials. Flow cytometry, nutrient, and temperature data are available in Dryad (https://doi.org/10.5061/dryad.q573n5tfr). Flowcam data is available on Ecotaxa under the project “Flowcam Composite Aquacosm_2018_VIMS-Ehux”. Sequencing data has been deposited under NCBI Bioproject PRJNA694552: 16S data is available under Biosample SAMN17576248 and 18S data is available under Biosample SAMN20295136. Additional data related to this paper may be requested from the authors.

## Notes

https://datadryad.org/stash/dataset/doi:10.5061/dryad.q573n5tfr

### Competing Interest Statement

The authors have declared no competing interest.

## References

1. Behrenfeld, M. J. et al. Climate-driven trends in contemporary ocean productivity. Nature 444, 752–755 (2006).

2. Schleyer, G. & Vardi, A. Algal blooms. Curr. Biol. 30, R1116–R1118 (2020).

3. Behrenfeld, M. & Boss, E. S. Resurrecting the Ecological Underpinnings of Ocean Plankton Blooms. Annu. Rev. Mar. Sci. i 6, 167–94 (2014).

4. Pomeroy, L. R. The Ocean’s Food Web, A Changing Paradigm. Bioscience 24, 499–504 (1974).

5. Azam, F. et al. The Ecological Role of Water-Column Microbes in the Sea. Mar. Ecol. Prog. Ser. 10, 257–263 (1983).

6. Wilhelm, S. W. & Suttle, C. A. Viruses and Nutrient Cycles in the Sea. Bioscience 49, 781–788 (1999).

7. Weinbauer, M. Ecology of prokaryotic viruses. FEMS Microbiol. Rev. 28, 127–181 (2004).

8. Guidi, L. et al. Plankton networks driving carbon export in the oligotrophic ocean. Nature 532, 465 (2016).

9. Vincent, F., Sheyn, U., Porat, Z., Schatz, D. & Vardi, A. Visualizing active viral infection reveals diverse cell fates in synchronized algal bloom demise. Proc. Natl. Acad. Sci. U. S. A. 118, e2021586118 (2021).

10. Dymond, J. & Lyle, M. Flux comparisons between sediments and sediment traps in the eastern tropical Pacific: Implications for atmospheric C02 variations during the Pleistocene. Limnol. Oceanogr. 30, 699–712 (1985).

11. Bratbak, G., Egge, J. K. & Heldal, M. Viral mortality of the marine alga Emiliania huxleyi (Haptophyceae) and termination of algal blooms. Mar. Ecol. Prog. Ser. 93, 39–48 (1993).

12. Wilson, W. H. et al. Isolation of viruses responsible for the demise of an Emiliania huxleyi bloom in the English Channel. J. Mar. Biol. Assoc. 82, 369–377 (2002).

13. Lehahn, Y. et al. Decoupling physical from biological processes to assess the impact of viruses on a mesoscale algal bloom. Curr. Biol. 24, 2041–2046 (2014).

14. Nejstgaard, J. C., Gismervik, I. & Solberg, P. T. Feeding and reproduction by Calanus finmarchicus, and microzooplankton grazing during mesocosm blooms of diatoms and the coccolithophore Emiliania huxleyi. Mar. Ecol. Prog. Ser. 147, 197–217 (1997).

15. Kalscheur, K., Rojas, M., Peterson, C., Kelly, J. & Gray, K. Algal exudates and stream organic matter influence the structure and function of denitrifying bacterial communities. Microb. Ecol. 64, 881–892 (2012).

16. Grossart, H.-P. & Simon, M. Interactions of planktonic algae and bacteria: effects on algal growth and organic matter dynamics. Aquat. Microb. Ecol. 47, 163–176 (2007).

17. Amin, S. A. et al. Interaction and signalling between a cosmopolitan phytoplankton and associated bacteria. Nature 522, 98–101 (2015).

18. Shibl, A. A. et al. Diatom modulation of select bacteria through use of two unique secondary metabolites. Proc. Natl. Acad. Sci. 117, 27445–27455 (2020).

19. Hahnke, R. et al. Dilution cultivation of marine heterotrophic bacteria abundant after a spring phytoplankton bloom in the North Sea. Environ. Microbiol. 17, 3515–3526 (2015).

20. Teeling, H. et al. Recurring patterns in bacterioplankton dynamics during coastal spring algae blooms. Elife e11888 (2016). doi:10.7554/eLife.11888.001

21. Giovannoni, S. J. SAR11 Bacteria: The Most Abundant Plankton in the Oceans. Ann. Rev. Mar. Sci. 9, 231–255 (2017).

22. Gonzalez, J. M. et al. Bacterial community structure associated with a dimethylsulfoniopropionate-producing North Atlantic algal bloom. Appl. Environ. Microbiol. 66, 4237–4246 (2000).

23. Keller, M. D. Biological Oceanography Dimethyl Sulfide Production and Marine Phytoplankton: The Importance of Species Composition and Cell Size. Biol. Oceanogr. 6, 375–382 (1988).

24. Barak-Gavish, N. et al. Bacterial virulence against an oceanic bloom-forming phytoplankter is mediated by algal DMSP. Sci. Adv. 4, eaau5716 (2018).

25. Tripp, H. J. et al. SAR11 marine bacteria require exogenous reduced sulphur for growth. Nature 452, 741–744 (2008).

26. Buchan, A., Lecleir, G. R., Gulvik, C. A. & González, J. M. Master recyclers: features and functions of bacteria associated with phytoplankton blooms. Nat. Rev. Microbiol. 12, 686–698 (2014).

27. Weinbauer, M. G. et al. Synechococcus growth in the ocean may depend on the lysis of heterotrophic bacteria. J. Plankton Res. 33, 1465–1476 (2011).

28. Schatz, D. et al. Ecological significance of extracellular vesicles in modulating host-virus interactions during algal blooms. ISME J. 1–8 (2021).

29. Weitz, J. S. et al. A multitrophic model to quantify the effects of marine viruses on microbial food webs and ecosystem processes. ISME J. 9, 1352–1364 (2015).

30. McManus, G. B. & Fuhrman, J. A. Control of marine bacterioplankton populations: Measurement and significance of grazing. Hydrobiologia 159, 51–62 (1988).

31. Cavalier-Smith, T. & Chao, E. E.-Y. Phylogeny and Megasystematics of Phagotrophic Heterokonts (Kingdom Chromista). J. Mol. Evol. 62, 388–420 (2006).

32. Bongiorni, L., Pusceddu, A. & Danovaro, R. Enzymatic activities of epiphytic and benthic thraustochytrids involved in organic matter degradation. Aquat. Microb. Ecol. 41, 299–305 (2005).

33. Poff, K. E., Leu, A. O., Eppley, J. M., Karl, D. M. & DeLong, E. F. Microbial dynamics of elevated carbon flux in the open ocean’s abyss. Proc. Natl. Acad. Sci. U. S. A. 118, e2018269118 (2021).

34. Raghukumar, S. The Marine Environment and the Role of Fungi. in Fungi in Coastal and Oceanic Marine Ecosystems 17–38 (Springer International Publishing, 2017).

35. Nakai, R. & Naganuma, T. Diversity and ecology of thraustochytrid protists in the marine environment. in Marine Protists: Diversity and Dynamics 331–346 (Springer Japan, 2015).

36. Bennett, R., Honda, D., Beakes, G. & Thines, M. Labryinthulomycota. in The handbook of theprotists (eds. JM, A., AGB, S. & Slamovits CH) 14:507–542 (Springer International Publishing, 2017).

37. Perkins, F. O. The ultrastructure of holdfasts, ‘rhizoids’, and ‘slime tracks’ in thraustochytriaceous fungi and Labyrinthula spp. Arch. Mikrobiol. 84, 95–118 (1972).

38. Hassett, B. T. A Widely Distributed Thraustochytrid Parasite of Diatoms Isolated from the Arctic Represents a gen. and sp. nov. J. Eukaryot. Microbiol. 67, 480–490 (2020).

39. Raghukumar, C. Thraustochytrid Fungi Associated with Marine Algae. Indian Jounral Mar. Sci. 15, 121–122 (1986).

40. Laundon, D., Mock, T., Wheeler, G. & Cunliffe, • Michael. Healthy herds in the phytoplankton: the benefit of selective parasitism. ISME J. 15, 2163–2166 (2021).

41. Kimura, H., Fukuba, T. & Naganuma, T. Biomass of thraustochytrid protoctists in coastal water. Mar. Ecol. Prog. Ser. 189, 27–33 (1999).

42. Ishibashi, Y., Aoki, K., Okino, N., Hayashi, M. & Ito, M. A thraustochytrid-specific lipase/phospholipase with unique positional specificity contributes to microbial competition and fatty acid acquisition from the environment. Sci. Reports 2019 91 9, 1–17 (2019).

43. Laber, C. P. et al. Coccolithovirus facilitation of carbon export in the North Atlantic. Nat. Microbiol. 3, 537–547 (2018).

44. Collins, J. R. et al. The multiple fates of sinking particles in the North Atlantic Ocean. Global Biogeochem. Cycles 29, 1471–1494 (2015).

45. Ramond, P. et al. Coupling between taxonomic and functional diversity in protistan coastal communities. Environ. Microbiol. 21, 730–749 (2019).

46. Holligan, P. M. et al. A biogeochemical study of the coccolithophore, Emiliania huxleyi, in the North Atlantic. Global Biogeochem. Cycles 7, 879–900 (1993).

47. Balch, W. M. et al. Factors regulating the Great Calcite Belt in the Southern Ocean and its biogeochemical significance. Global Biogeochem. Cycles 30, 1124–1144 (2016).

48. Alldredge, A., Passow, U. & Haddock, S. H. D. The characteristics and transparent exopolymer particle (TEP) content of marine snow formed from thecate dinoflagellates. J. Plankton Res. 20, 393–397 (1998).

49. Passow, U. et al. The origin of transparent exopolymer particles (TEP) and their role in the sedimentation of particulate matter. Cont. Shelf Res. 21, 327–346 (2001).

50. Vidal-Melgosa, S. et al. Diatom fucan polysaccharide precipitates carbon during algal blooms. Nat. Commun. 2021 121 12, 1–13 (2021).

51. Vidal-Melgosa, S. et al. A new versatile microarray-based method for high throughput screening of carbohydrate-active enzymes. J. Biol. Chem. 290, 9020–9036 (2015).

52. Torode, T. A. et al. Dynamics of cell wall assembly during early embryogenesis in the brown alga Fucus. J. Exp. Bot. 67, 6089–6100 (2016).

53. Torode, T. A. et al. Monoclonal Antibodies Directed to Fucoidan Preparations from Brown Algae. PLoS One 10, e0118366 (2015).

54. Damare, V. & Raghukumar, S. Marine aggregates and transparent exopolymeric particles (TEPs) as substrates for the stramenopilan fungi, the thraustochytrids: Roller table experimental approach. Kavaka 40, 22–31 (2012).

55. Vardi, A. et al. Host – virus dynamics and subcellular controls of cell fate in a natural coccolithophore population. Proc. Natl. Acad. Sci. U. S. A. 109, 19327–19332 (2012).

56. Blanco-Ameisjeira, S. et al. Phenotypic variability in the coccolithophore Emiliania huxleyi. PLoS One 11, 1–17 (2016).

57. Forterre, P. The virocell concept and environmental microbiology. ISME J. 7, 233–236 (2013).

58. Rosenwasser, S., Ziv, C., Graff Van Creveld, S. & Vardi, A. Virocell Metabolism: Metabolic Innovations During Host-Virus Interactions in the Ocean. Trends Microbiol. 24, 821–832 (2016).

59. Emerson, S. & Hedges, J. Chemical oceanography and the marine carbon cycle. in Chemical Oceanography and the Marine Carbon Cycle 1–461 (Cambridge University Press, 2008).

60. Blanco-Ameijeiras, S. et al. Phenotypic Variability in the Coccolithophore Emiliania huxleyi. PLoS One 11, e0157697 (2016).

61. Klawonn, I. et al. Characterizing the “fungal shunt”: Parasitic fungi on diatoms affect carbon flow and bacterial communities in aquatic microbial food webs. Proc. Natl. Acad. Sci. USA 118, (2021).

62. Kimura, H. & Naganuma, T. Thraustochytrids: A neglected agent of the marine microbial food chain. Aquat. Ecosyst. Health Manag. 4, 13–18 (2001).

63. Kuhlisch, C. et al. Viral infection of algal blooms leaves a unique metabolic footprint on the dissolved organic matter in the ocean. Sci. Adv. 7, eabf4680 (2021).

64. Sheyn, U. et al. Expression profiling of host and virus during a coccolithophore bloom provides insights into the role of viral infection in promoting carbon export. ISME J. 12, 704–713 (2018).

65. Pais-Correia, A. M. et al. Biofilm-like extracellular viral assemblies mediate HTLV-1 cell-to-cell transmission at virological synapses. Nat. Med. 16, 83–89 (2010).

66. Thoulouze, M. I. & Alcover, A. Can viruses form biofilms? Trends in Microbiology 19, 257–262 (2011).

67. Johns, C. T. et al. The mutual interplay between calcification and coccolithovirus infection. Environ. Microbiol. 21, 1896–1915 (2019).

