## Supplementary Figures for "Viral infection switches the balance between bacterial and eukaryotic recyclers of organic matter during algal blooms"

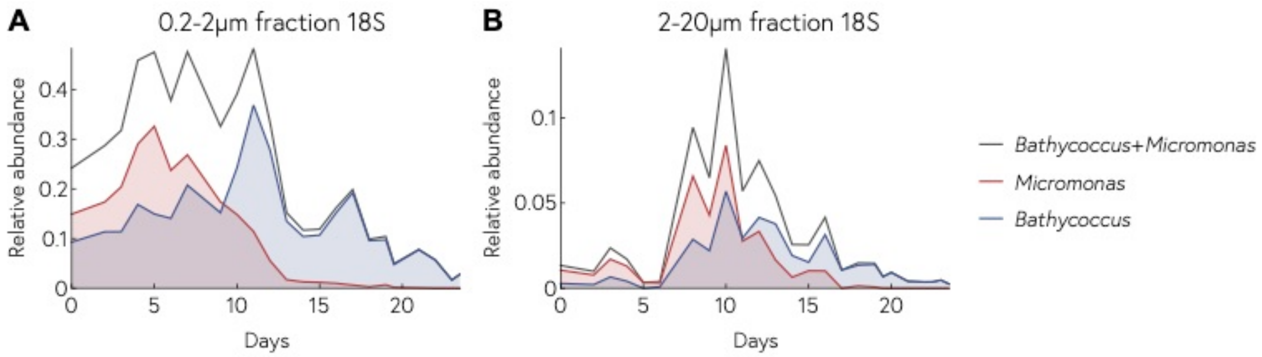


**Supplementary Figure 1**: *Bathycoccus* and *Micromonas* relative abundances in (A) the 0.2-2µm fraction and (B) 2-20µm fraction of the 18S microbiome. The relative abundances of *Micromonas* (red) and *Bathycoccus* (blue) are summed in black and represent over 40% of all the reads in the 0.2-2µm size fraction.

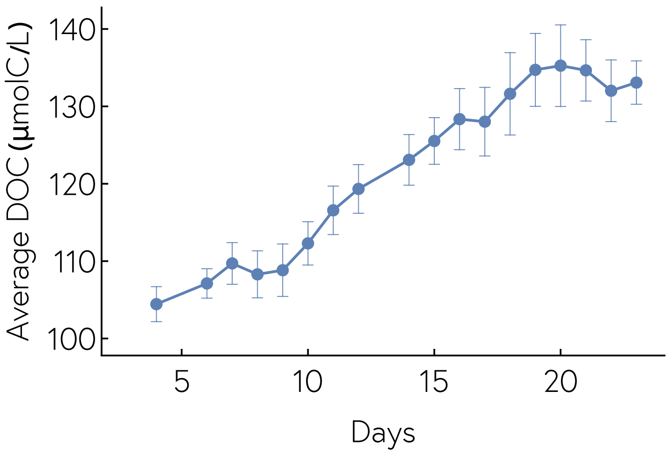


**Supplementary Figure 2:** Dissolved organic carbon (DOC) concentration through time. Concentrations are averaged across all bags and smoothed across three days with bars representing standard deviation.

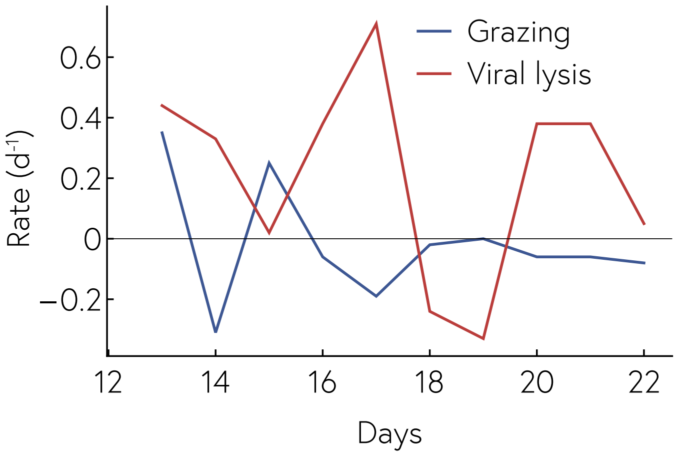


**Supplementary Figure 3:** Rates of grazing (blue) and viral lysis (red) measured by paired dilution assay through time, using a mixture of water from bags 1-4. See Methods for calculation of rates.

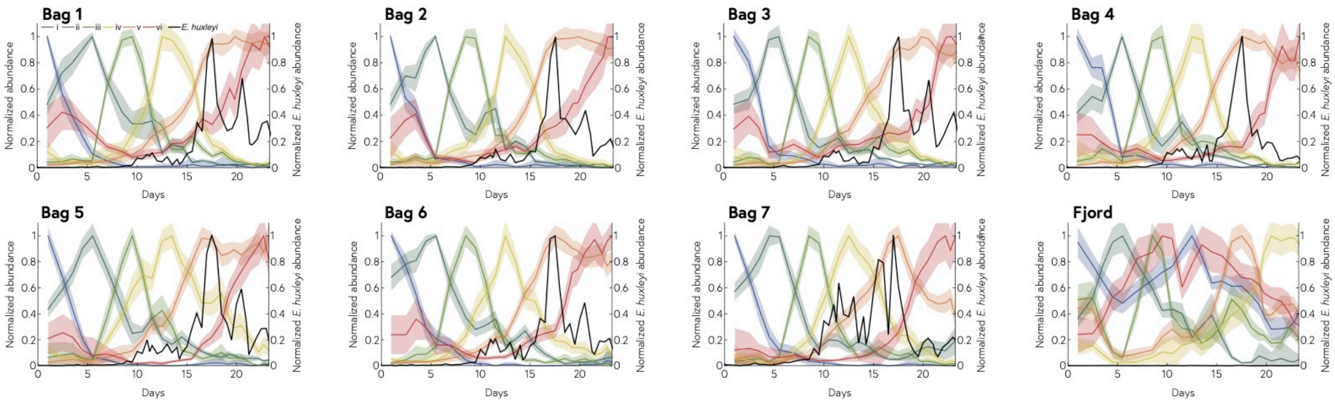


**Supplementary Figure 4:** Succession of eukaryotic species within each bag normalized per cluster. Species are clustered by similarity of their relative abundance dynamics. The shaded area represents the standard deviation within each cluster. Each cluster is normalized to its own maximum abundance and clusters composition is detailed Fig. 2a. *E. huxleyi* abundance is overlaid in black and normalized to each bag's maximum value.

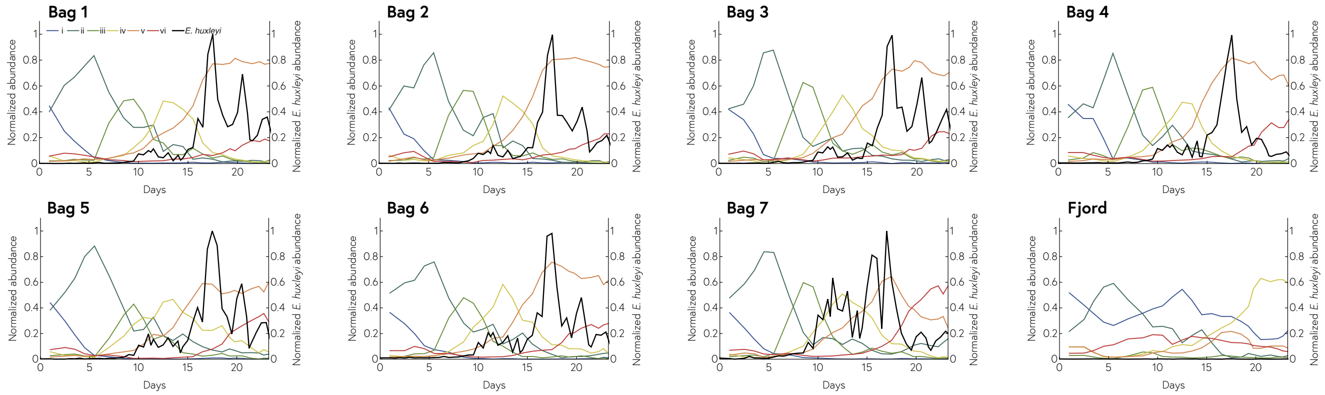


**Supplementary Figure 5:** Succession of eukaryotic clusters within each bag normalized to the total relative abundance. Species are clustered by similarity of their relative abundance dynamics. Each cluster is normalized to the total relative abundance and their species composition is detailed Fig. 2a. *E. huxleyi* abundance is overlaid in black and normalized to each bag's maximum value.

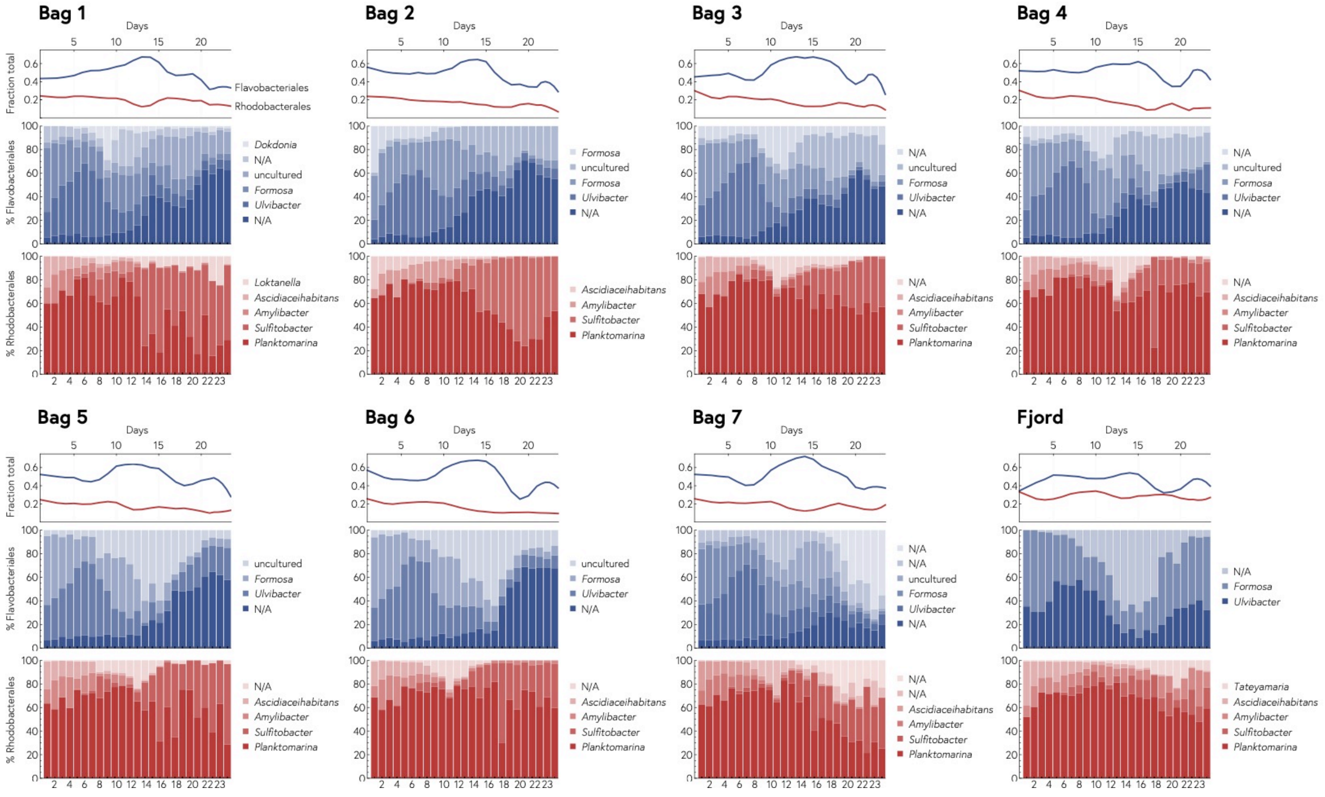


**Supplementary Figure 6:** Flavobacteriales (blue) and Rhodobacterales (red) species relative abundances over time in each bag. The top pannel represents the total relative abundance of Flavobacteriales (blue) and Rhodobacterales (red) species within the whole bacterial community.

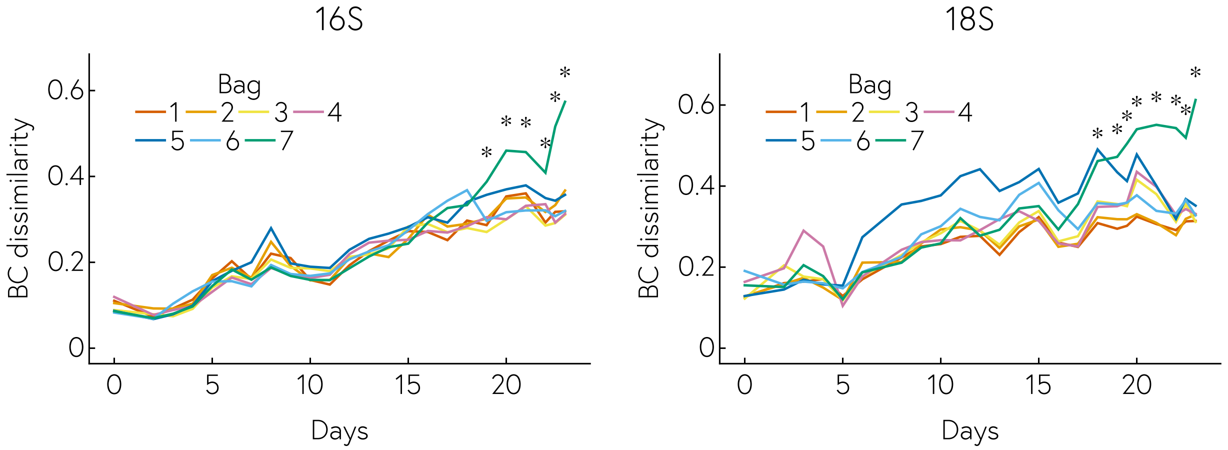


**Supplementary Figure 7:** Divergence of bacterial (16S) and nanoeukaryotic (18S) compositions between bags over time. For the 18S and 16S separately, a Bray-Curtis distance between the bags was measured for each day. The stars indicate significant differences between bag 7 and the other bags according a Kolmogorov-Smirnov test with Benjamini-Hochberg multiple testing correction.

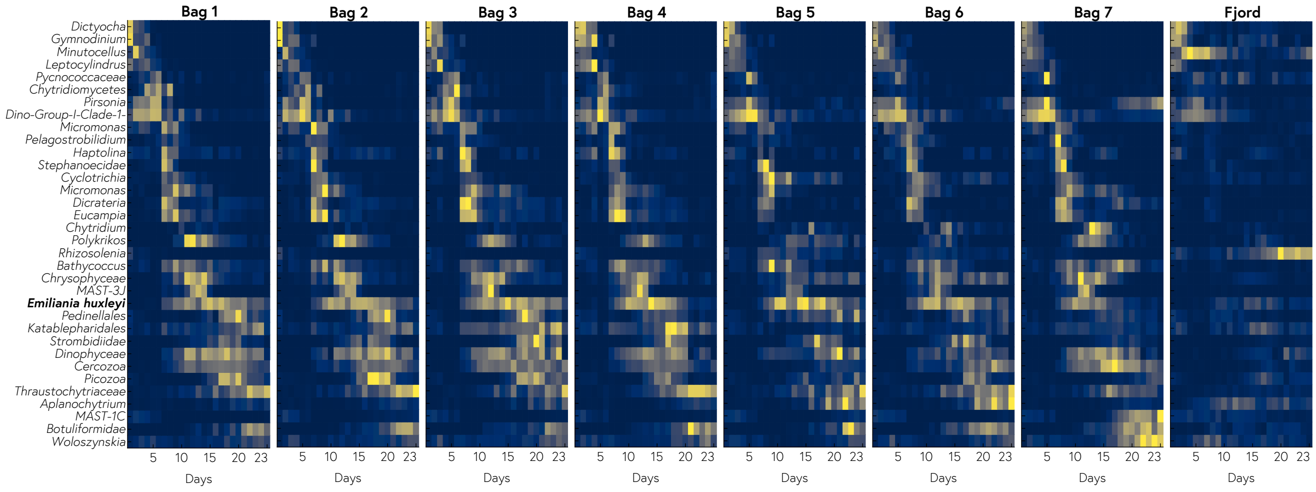


**Supplementary Figure 8:** Most abundant 18S ASVs in all bags in the 2µm size fraction. The heatmap is normalized per row, meaning each ASV is normalized to its maximum abundance across all bags and all days.

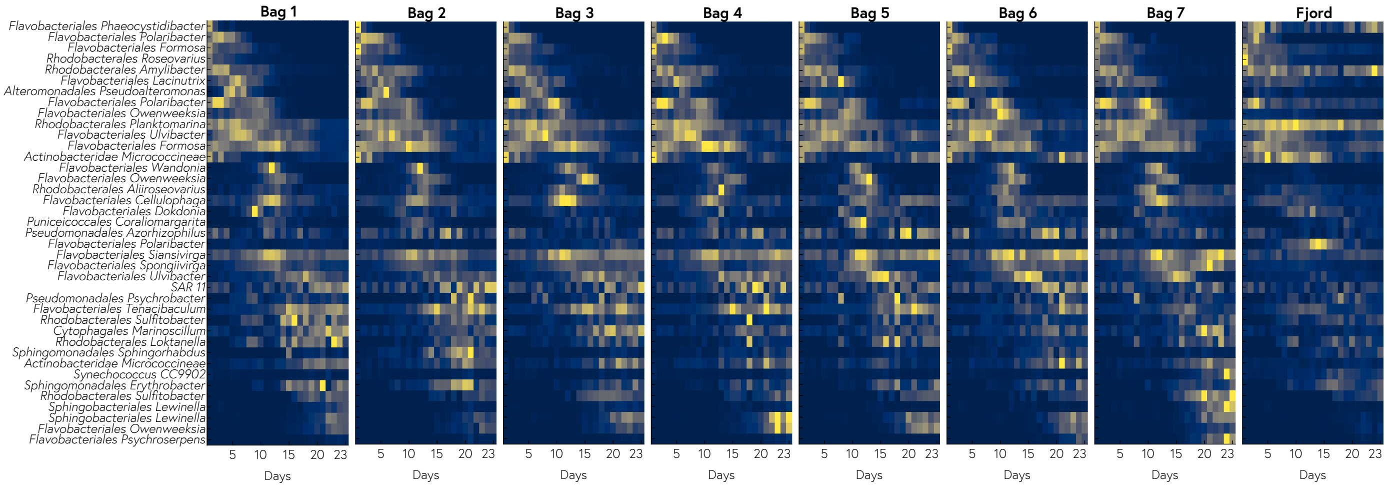


**Supplementary Figure 9:** Most abundant 16S ASVs in all bags in the 0.2μm size fraction. The heatmap is normalized per row, meaning each ASV is normalized to its maximum abundance across all bags and all days.

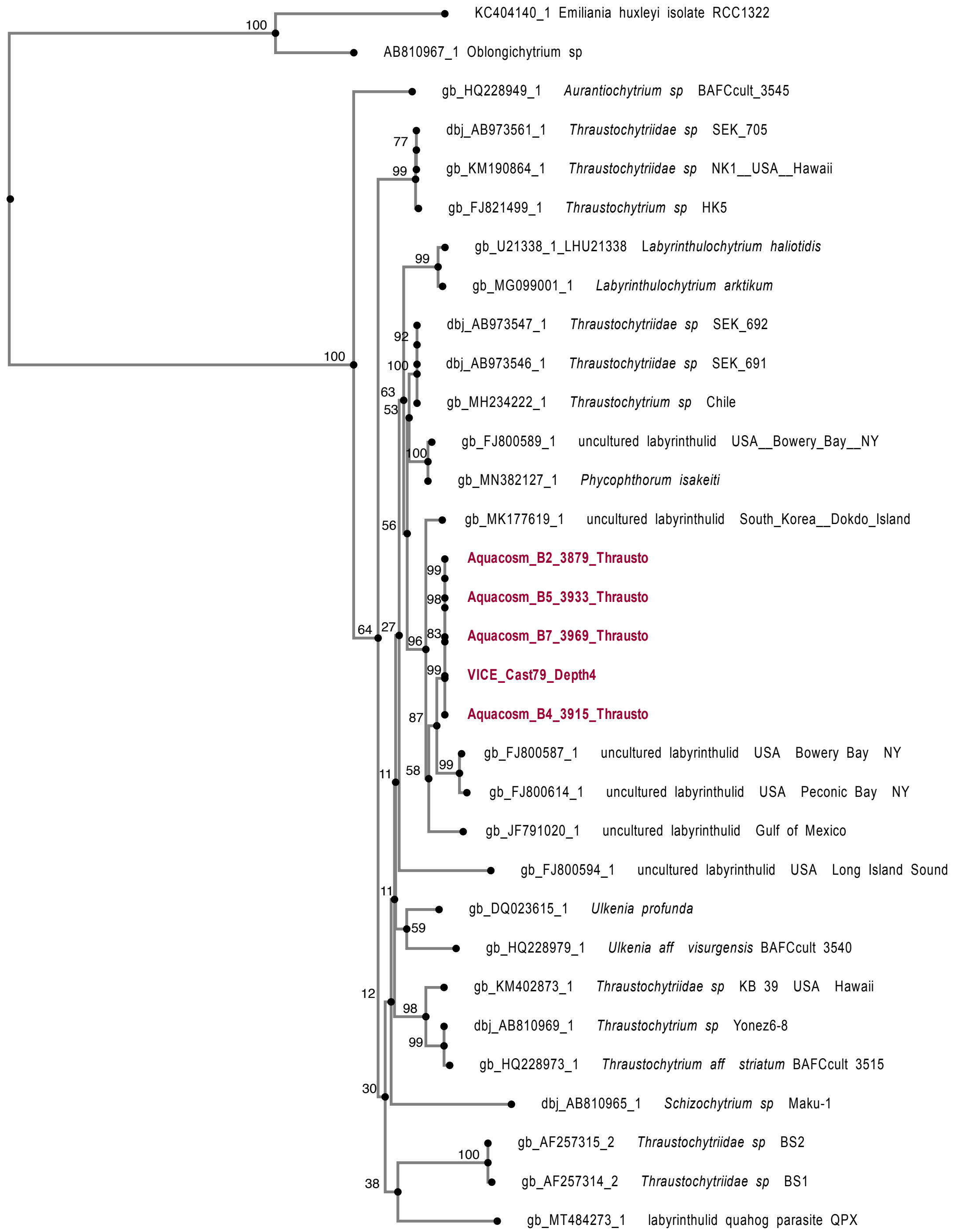


**Supplementary Figure 10:** Phylogeny of thraustochytrids including new sequences from this study (in red). Aquacosm sequences all come from the Norwegian fjord. The VICE sequence comes from samples collected in the North Atlantic. The tree was based on 866 conserved sites and numeric values on the nodes were obtained with 1000 bootstraps. *E. huxleyi* and *Oblongichytrium* were used as outgroups (see Methods).

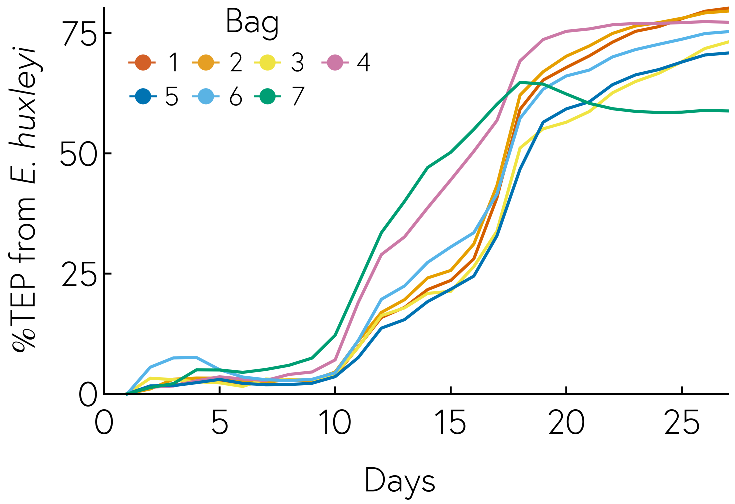


**Supplementary Figure 11:** Predicted fraction of transparent exopolymers (TEP) contributed by *E. huxleyi* for each mesocosm enclosure. TEP was modelled as the sum of *E. huxleyi,* naked nanophytoplankton, and picophytoplankton TEP production, with a loss factor corresponding to TEP degradation or sinking (see Methods).

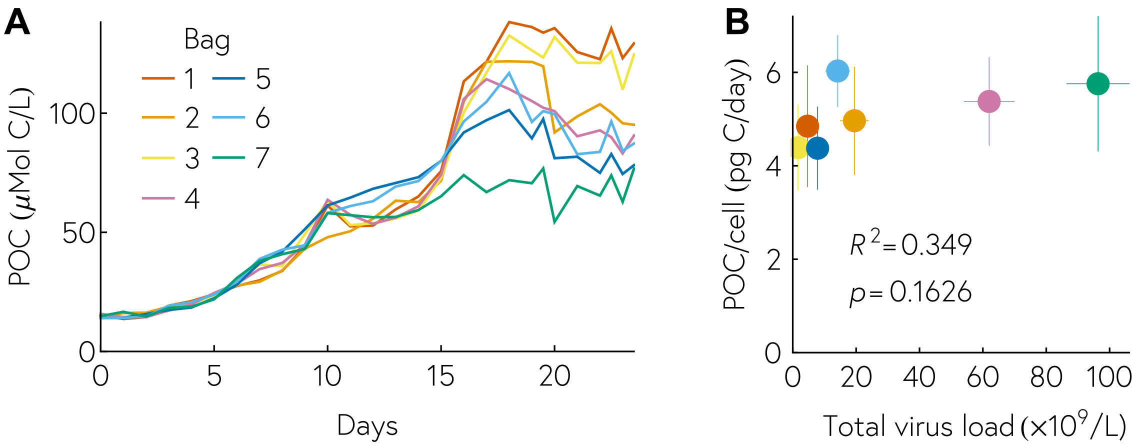


**Supplementary Figure 12:** Measurements and modelling of particulate organic carbon (POC). (A) POC measurements over time for each individual mesocosm enclosure. (B) Predicted POC/cell for each individual enclosure, as a function of total viral load in each bag. The low $R^2$ and *p* values show that POC measurements cannot be well predicted by our model, contrary to TEP and PIC.

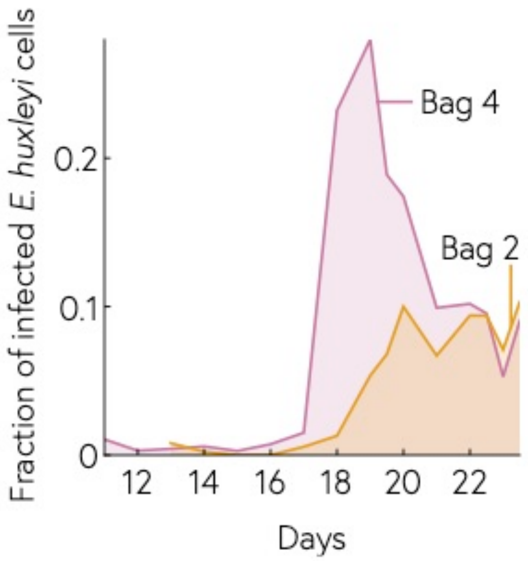


**Supplementary Figure 13:** Fraction of infected *E. huxleyi* cells, measured by smFISH in bags 2 and 4. After fixation, cells were stained with a 28S probe designed to detect *E. huxleyi* cells and a probe targeting the viral *mcp* mRNA to identify actively infected cells.

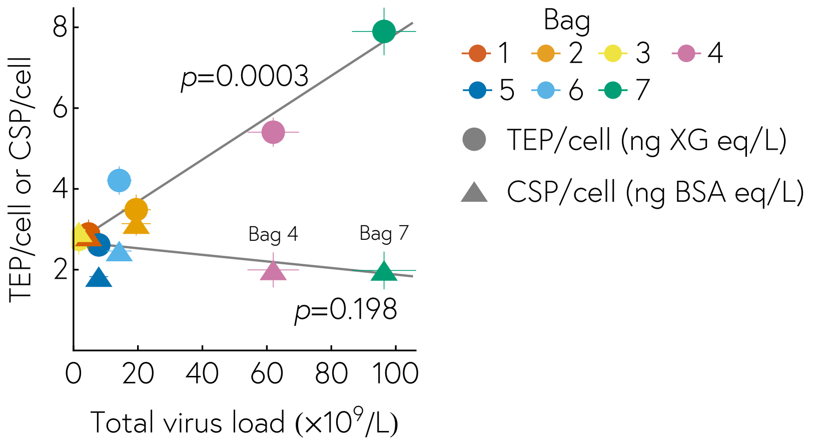


**Supplementary Figure 14:** Comparison of transparent exopolymer particles (TEP) and Coomassie stainable particles (CSP) modelling. Predicted TEP and CSP per cell as a function of viral load using the same model. Non-significant fits for the CSP modelling shows that our model cannot explain CSP, contrary to TEP and PIC.
