## Supplementary Methods for "Viral infection switches the balance between bacterial and eukaryotic recyclers of organic matter during algal blooms"

***Methods for data analysis in figures***
All analyses in figures were performed using the Mathematica 12.3 software (Wolfram, Champaign, USA).

***Fig. 1***
C&D. To calculate integrated abundances of *E. huxleyi* cells and EhV, we first selected days for which all the bags had a non-null value. Values were then summed up to obtain the integrated abundance.

E&J. To correct for enclosure cover, which systematically lowered *E. huxleyi* abundance, we added the average difference between *E. huxleyi* abundance in covered and uncovered bags to the *E. huxleyi* abundance in each uncovered bag. This is equivalent to fitting total *E. huxleyi* abundance with a vector encoding the bag cover and then keeping only the fit residuals. We then computed a standard linear fit between the corrected *E. huxleyi* total abundances and total EhV abundances. We followed the same procedure for the correlations in panel J.

***Fig. 2***
A. The ASVs that were selected appeared at a relative abundance of at least 2% in at least 4 samples for the 0.2-2µm 16S sequences and at least in 8 samples for the 2-20µm 18S sequences. Abundances were concatenated for each time point and normalized by row, to have maximum relative abundance of 1 across all samples. ASVs were sorted by the position of their individual center of mass $t_{CM}$ defined by $t_{CM} = \frac{\sum_i t_i f(t_i)}{\sum_i f(t_i)}$ with *i* representing the different time points and $f(t_i)$ the relative abundance of the ASV. The same figure for the individual bags in shown in **Supplementary Fig. 8, 9**.

B. We selected 18S ASVs with a maximum relative abundance of at least 2% and observed in at least 5 samples. We averaged relative abundance across bags and then smoothed the time series with a moving average filter (width 2). Then, we grouped all ASVs into clusters based on their cosine distance using Mathematica's FindClusters function and the KMeans method. The number of possible clusters ranged from 2 to 12, and the final number of clusters was decided using the silhouette method[1]. Only silhouette scores for 2 and 6 clusters were positive (between-cluster distance minus within-cluster distance).

D. We subset reads that map to either flavobacteriales or rhodobacterales, then renormalized within each class, taking the mean over bags. Results per bag are shown in **Supplementary Fig. 6**.

E. The turnover time was defined by the exponential rate *k* at which the Bray-Curtis similarity $BC(t)$ declined over time. To this end, for a given bag, we computed the Bray-Curtis similarity between the composition vector at a starting day $t'$ with all following days *t*, giving a curve that declined roughly exponentially. For earlier starting days (for which the similarity curves declined the furthest), we found that the Bray-Curtis similarity never reached 0 but instead leveled out around $BC_\infty = 0.05$ (due to ASVs that are constantly present in all the samples and maintain a minimal level of similarity between bags). Thus, we imposed an offset at $BC_\infty$ for all fits (using Mathematica's FindFit function) with the function:

$$BC(t) = (1 - BC_\infty) * e^{-k(t'-t)} + BC_\infty$$

The turnover is averaged over bags, showing the standard deviation as error bars in the figure.

F. The divergence between bags has been calculated as follows: we first measured, for each bag, the Bray-Curtis distance between this given bag and all the other bags at the end of the experiment (**Supplementary Fig. 7**). In order to control for the existing differences between bags at the beginning of the bloom, Bray-Curtis distances were normalized according to the differences between bags at the starting day of the *E. huxleyi* bloom. As the exact starting days of the bloom is not clear, we normalized for starting days 11, 12, or 13. The plot shows averages with the standard deviation as error bars. For the 18S microbiome, we first removed reads that map to *E. huxleyi* to reduce bias towards bag 7 (which has by far the lowest *E. huxleyi* abundance, **Fig. 1c**). Panel H contains the same data but plotted against the total viral abundance in each bag.

G. We found 16S and 18S ASVs that were different in bag 7 by manual inspection of each ASV's relative abundance between bags. Then, we performed a Kolmogorov-Smirnov test between the distances for those ASV within bags 1-6 and those between bags 1-6 and bag 7. From the most differentially abundant ASVs we selected four to be shown in panel G.

***Fig 3.***

A. Functional annotation of dominant 18S ASVs was based on manual literature search for the 100 most abundant 18S ASVs. Automatic annotation using the functional database created by[2] gave qualitatively identical results but contained fewer organisms (covering about 50% of reads). The relative abundance of each trait was obtained by summing up the relative abundance of all the species harboring a specific trait. We used the annotations from[2] to further subdivide heterotrophs into osmotrophs, saprotrophs, and other types of heterotrophy (e.g., grazing), ignoring ASVs with missing annotations.

D. To quantify the rate of change *k* of the biomass ratio of thraustochytrids to bacteria we fit a linear function to the log of biomass ratio from day 10 to the time point *t* where the ratio was maximal; for bag 7, this was day 18, for all others, day 23. We thus have:

$$\log BR(t) = kt + \log BR(0)$$

***Fig. 4.***

C&D. Since TEP accumulates over time, it cannot be expressed as a weighted sum of phytoplankton abundances. Instead, we formulate the model as a recursive relation where TEP can be produced by *E. huxleyi*, naked nanophytoplankton, and picophytoplankton, and degraded or lost through sinking:

$$TEP(t) = (1-d)TEP(t-1) + a_E E(t) + a_N N(t) + a_P P(t),$$

The amount of TEP at time $t$ is given by the fraction (1-$d$) of TEP at time $t$-1, where $d$ corresponds to the fraction of TEP that is degraded between time points, plus the amount of TEP produced by the phytoplankton cells present at time $t$ (or time $t$-1, which gives equivalent results). E, N, and P correspond to *E. huxleyi*, naked nanophytoplankton, and picophytoplankton, respectively. The parameter $a_E$ corresponds to the amount of TEP produced per *E. huxleyi* cell, reported in panel D. $a_E$ is set to be fixed through time, and different for each bag. This recursion can be solved to give an explicit expression for TEP(t):

$$TEP(t) = \sum_{t'=0}^{t} (1-d)^{t-t'} [a_E E(t') + a_N N(t') + a_P P(t')].$$

This functional form was then used to perform a linear model fitting with the constraint $a_i \geq 0$ for various values of the parameter *d*. The best fit, defined by maximum $R^2$ over the resulting linear model, was used to fix *d*=0.12.

E. Using the smFISH method that reports the proportion of infected *E. huxleyi* cells, we estimated the amount of TEP produced from infected cells. We first used the least infected uncovered bags (bags 1 and 3) as a baseline to fix model parameters such as how much TEP does a non-infected cell produce. We then split the *E. huxleyi* abundance into an uninfected subpopulation producing $T$ TEP/cell as in the uninfected bags, and an infected subpopulation producing $I*T$ TEP/cells. To define I, we combined the fixed model parameters (i.e., amount of TEP produced per cell from **Fig. 4d** for bags 1 and 3) with the measured fraction of infected cells. We adjusted the factor $I=4.5$ to minimize deviation of the measure total TEP concentration from the model prediction including the two subpopulations. The same procedure was used for panel H, using the corresponding model for PIC.

F&G. To model the amount of PIC produced per cell we assume that the measured PIC only increases via new *E. huxleyi* coccoliths. The equivalent model for PIC reads

$$PIC(t) = (1-d)PIC(t-1) + a_E \max\big(E(t) - E(t-1)\big).$$

Where $a_E$ is the amount of PIC produced per cell, and displayed in panel G. Using the same procedure as for TEP, we obtain the best fit for *d*=0.0075.

### *Methods for data collection*

***Mesocosm core setup***. The mesocosm experiment AQUACOSM VIMS-Ehux was carried out for 24 days between 24th May (day 0) and 16th June (day 23) 2018 in Raunefjorden at the University of Bergen's Marine Biological Station Espegrend, Norway (60°16′11N; 5°13′07E). The experiment consisted of seven enclosure bags made of transparent polyethylene (11 $m^3$, 4 m deep and 2 m wide, permeable to 90% photosynthetically active radiation) mounted on floating frames and moored to a raft in the middle of the fjord. The bags were filled with surrounding fjord water (day -1; pumped from 5 m depth) and continuously mixed by aeration (from day 0 onwards). Each bag was supplemented with nutrients at a nitrogen to phosphorus ratio of 16:1 (1.6 µM $NaNO_3$ and 0.1 µM $KH_2PO_4$ final concentration) on days 0-5 and 14-17, whereas on days 6, 7 and 13 only nitrogen was added.

***Measurement of dissolved inorganic nutrients.*** Unfiltered seawater aliquots (10 mL) were collected from each bag and the surrounding fjord water in 12 mL polypropylene tubes and stored frozen at −20 °C. Dissolved inorganic nutrients were measured with standard segmented flow analysis with colorimetric detection[3], using a Bran & Luebe autoanalyser. Data is available in [4].

***Measurement of water temperature and salinity.*** Water temperature and salinity were measured in each bag and the surrounding fjord water using a SD204 CTD/STD (SAIV A/S, Laksevag, Norway). Data points were averaged for 1-3 m depth (descending only). When this depth was not available, the available data points were taken. Data is missing for the fjord in days 0-1. Outliers were removed for the following samples: bag 1 at days 0, 4, 15; bag 7 at day 15. Data is available in[4].

***Flow Cytometry measurements.*** Samples for flow cytometric counts were collected twice a day, in the morning (7:00 a.m.) and evening (8:00-9:00 p.m.) from each bag and the surrounding fjord, which served as an environmental reference. Water samples were collected in 50 mL centrifugal tubes from 1 m depth, pre-filtered using 40 µm cell strainers, and immediately analyzed with an Eclipse iCyt (Sony Biotechology, Champaign, IL, USA) flow cytometer. A total volume of 300 µL with a flow rate of 150 µL/min was analyzed. A threshold was applied based on the forward scatter signal to reduce the background noise.
Phytoplankton populations were identified by plotting the autofluorescence of chlorophyll versus phycoerythrin and side scatter: calcified *E. huxleyi* (high side scatter and high chlorophyll), *Synechococcus* (high phycoerythrin and low chlorophyll), nano- and picophytoplankton (high and low chlorophyll, respectively). Chlorophyll fluorescence was detected by FL4 (excitation (ex): 488nm and emission (em): 663-737 nm). Phycoerythrin was detected by FL3 (ex: 488 nm and em: 570-620 nm). Raw .fcs files were extracted and analyzed in R using 'flowCore' and 'ggcyto' packages.
For bacteria counts, 200 µL of sample were fixed with 4 µL of 20% glutaraldehyde (final concentration of 0.5%) for one hour at 4°C and flash frozen. They were thawed and stained with SYBR gold (Invitrogen) that was diluted 1:10,000 in Tris-EDTA buffer, incubated for 20 min at 80°C and cooled to room temperature. Bacteria were counted and analyzed using a Cytoflex and identified based on the Violet SSC-A versus FITC-A by comparing to reference samples containing fixed bacteria from lab cultures. A total volume of 60 µL with a flow rate of 10 µL/min was analyzed. A threshold was applied based on the forward scatter signal to reduce the background noise. For plotting bacteria (**Fig. 1h**), a sliding window of size 3 was used. Data is available in[4].

***Enumeration of extracellular EhV abundance by qPCR.***
DNA extracts from filters from the core sampling (see above) were diluted 100 times, and 1 µL was then used for qPCR analysis. EhV abundance was determined by qPCR for the major capsid protein (*mcp*) gene: 5′-acgcaccctcaatgtatggaagg-3′ (mcp1F) and 5′-rtscrgccaactcagcagtcgt -3′ (mcp94Rv). All reactions were carried out in technical triplicates. For all reactions, Platinum SYBER Green qPCR SuperMix-UDG with ROX (Invitrogen, Carlsbad, CA, USA) was used as described by the manufacturer. Reactions were performed on a QuantStudio 5 Real-Time PCR System equipped with the QuantStudio Design and Analysis Software version 1.5.1 (Applied Biosystems, Foster City, CA, USA) as follows: 50°C for 2 min, 95°C for 5 min, 40 cycles of 95°C for 15 s, and 60° C for 30 s. Results were calibrated against serial dilutions of EhV201 DNA at known concentrations, enabling exact enumeration of viruses. Samples showing multiple peaks in melting curve analysis or peaks that were not corresponding to the standard curves were omitted.
Data is available in[4].

***FlowCam analysis.*** Samples for automated flow imaging microcopy were collected once a day in the morning (7:00 a.m.) from each bag and the surrounding fjord, which served as an environmental reference. Water samples were collected in 50 mL centrifugal tubes from 1 m depth, kept at 12°C in darkness, and analyzed within two hours of sampling, using a FlowCAM II (Fluid Imaging Technologies Inc., Scarborough, ME, USA) fitted with a 300 µm path length flow cell and a 4× microscope objective. Images were collected using auto-image mode at a rate of 7 frames/second. A sample volume of 10 mL was processed at a flow rate of 0.7 mL/min. Individual objects within each sample were clustered and annotated using the Ecotaxa platform[5]. Absolute counts for major groups, including the most abundant ciliate category Ciliophora U04, were then exported and normalized by the individual amount of water volume processed for each sample.

Data is available under "Flowcam Composite Aquacosm_2018_VIMS-Ehux" project on Ecotaxa.

***Core microbiome harvesting, sequencing, and annotation.*** Every day, between 1 to 2 L of water samples of each bag and fjord water were pre-filtered at 200 μm, then filtered sequentially through 20 μm and 2 μm, and finally 1-2 L filtrate was filtered through 0.22 μm hydrophilic polycarbonate filters (Isopore, 47 mm; Merck Millipore, Cork, Ireland). Filters were immediately flash frozen in liquid nitrogen and stored at -80°C until further processing. DNA was extracted from the 2 μm and 20 μm filters using the DNeasy PowerWater kit (Qiagen, Hilden, Germany) according to the manufacturer's instructions. 0.2 μm filters were extracted using DNAdvance Kit (Beckman-Coulter, Brea, USA).
The bacterial community was sequenced using the EMP 16S amplicon protocol and 515F-806R primers[6] at the Environmental Sample Preparation and Sequencing Facility (ESPSF), which is located in the Argonne National Laboratory. Degeneracy was added to the 515F primer to reduce bias against Crenarchaeota/Thaumarchaeota (also called 515F-Y[7]) and to the 806R primer to minimize the bias against the SAR11 clade (806R[8]). The primer sequences without the linker, pad, barcode, or adapter are as follows: 5' - GTGYCAGCMGCCGCGGTAA - 3'(515F-Y) 5' - GGACTACNVGGGTWTCTAAT - 3' (806R). ASVs were called using DADA2[9] with standard parameters. Taxonomic identity mapping was performed using RDP classifier[10]. The average sequencing depth per sample was about 27000 +/- 6200 (min 5500, max 49500 reads). Reads were normalized to the total amount of reads within each sample to convert them into relative abundance. Prior to further analysis, all reads that map to chloroplasts were removed (corresponding to up to 5% of all reads during the first bloom and 15% of all reads during the *E. huxleyi* bloom).
For the 18S sequencing of the DNA extracts from the 0.2 μm and 2 μm filters, the V4 region of the 18S rDNA sequence was amplified using the TAREuk454FWD1 (5′-CCAGCA(G/C)C(C/T)GCGGTAATTCC-3′) from[11] and a modified V4Rev_Piredda (5' - ACTTTCGTTCTTGATYRATGA - 3') from[12] in order to identify *E. huxleyi*, combined with CS1 and CS2 Illumina adaptors. We used the following PCR mix: 12.5 μL of Buffer myTAQ HS 2X Mix, 1 μL of each primer 0.4 μM final concentration, 0.75 μL of DMSO 3%, 8.75 μL of ultrapure water, 1 μL of DNA template. We used the following PCR conditions: initial denaturation of 2 min at 95°C followed by 10 cycles of 10 sec 95°C, 30 sec 53°C, 30 sec 72°C then 15 cycles of 10 sec 95°C, 30 sec 48°C, 30 sec 72°C, final elongation of 10 minutes at 72°C. PCR products were prepared for Illumina sequencing on a MiSeq 2x250. Fastq files were then cleaned and amplicon sequencing variants determined using the DADA2 pipeline[9], annotated with the PR2 database[13] and analyzed using the "phyloseq" package in R[14].
Data has been deposited under NCBI Bioproject PRJNA694552: 16S data is available under Biosample SAMN17576248 and 18S data is available under Biosample SAMN20295136.

***ddPCR quantification***.
*Thraustochytrids:* Digital droplet PCR (Bio-Rad, Hercules, USA) was performed on 2 μm mesocosm filters of days 2, 8, 14, 16, 18, 20, 23 of each bag including the fjord, to assess the absolute concentration of thraustochytrids. For VICE-cruise samples[15], representative samples of each bloom phase were chosen.
Primers targeting the 18S rDNA gene of thraustochytriaceae were used[16] with forward primer SYBR-ThF 5'-GGATCGAAGATGATTAGATACCA-3' and reverse primer SYBR-ThR 5'-GACTTTGATTTCTCATGTGC -3'. Primers were checked for specificity in PR2[13]. Sample mix consisted of 10 μL of 2X QX200 ddPCR EvaGreen supermix, 1 μL of 2uM forward primer, 1 μL of 2 μM reverse primer, 5 μL of water and 5 μL of the DNA sample. To load the optimal amount of DNA, DNA extractions were diluted 1:10 and DNA concentration was

measured using a Qubit dsDNA HS Assay Kit (Invitrogen, Waltham, USA). Depending on the concentration, between 1-5 µL of extracts were completed to a total of 5 µL with ultra-pure water, and used in the final ddPCR reaction. Less than 80 ng of DNA was used for each reaction. From the final mix of 22 µL, 20 µL of each sample were loaded in the DG8 Cartridge and inserted in the QX200 droplet generator. Each cartridge contained a negative control containing the ddPCR mix with 5 µL of water. After droplet generation, samples were transferred to a 96 well-plate and inserted in a C1000 Touch thermal cycler. The following cycle was used: 95°C 5 min, followed by 40 cycles of 96°C for 30 sec, 58°C for 1 min, 4°C 5 min, 90°C 5 min and infinite hold at 4°C. After thermal cycling, the 96-well plate was read in the QX200 Droplet Reader and results analyzed using the Quantasoft software.

Quantasoft provides a final concentration of target copies/µL of ddPCR reaction. For mesocosm samples, we first calculated the total amount of target copies in 20 µL of ddPCR reaction and normalized it by the amount of sea water that was sampled, to obtain a final concentration of target copies/mL of sampled sea water. To convert 18S copies/mL into cell/ml, we estimated the amount of 18S copies per thraustochytrid cell. The number of 18S rDNA copies/cell was calculated based on the relationship between genome size and copy number recently published in[17]. Published thraustochytrid genomes range between 38.7 Mb[18] and 43 Mb[19]. Using the regression equation on log transformed data with an average thraustochytrid genome size of 40 Mb, we obtain $f(x) = 0.6607(\log(40)) + 0.7508 = 1.809$ with $f(x)$ the log value of total 18S copies. We therefore obtain that the estimated 18S copy number in thraustochytrids cells is $10^{\wedge 1.809} = 64$ copies. The thraustochytrid biomass was estimated based on a value of $1.65 \times 10^{-10}$ g of C/cell[20]. The bacterioplankton biomass was estimated based on a value of $10 \times 10^{-15}$ g C/cell[21], using abundance counts from the flow cytometer. A detailed calculation for each sample is available in **Supplementary Table 1.** For cruise samples, we report copies per ng of extracted DNA.

***Sanger sequencing of thraustochytrids from environmental samples***

To identify thraustochytrid species from the mesocosm, DNA extracts from June 16th 2018 (Day 23) of the 2-20 µm size fraction from bag 2, bag 4, bag 5, and bag 7 were used. To identify thraustochytrids from an open ocean bloom, DNA extracts from the NA-VICE Cruise Cast 79[15], 28m depth was chosen for its high concentration of thraustochytrids based on ddPCR.

DNA from each sample was used as a template in PCR reactions with the primer 18S-F[22] and LABY-Y[23] (~1400 bp product). PCR reactions were made with Platinum Taq DNA Polymerase reagents (Invitrogen, Waltham, USA) as follows: 5 µl 10× Platinium Taq buffer, 1 µl 10 mM dNTPs, 1 µl 10 µM of forward and reverse primers, 1.5 µl 50 mM $MgCl_2$, 38.3 µl water; 0.2 µl Platinum Taq polymerase; and 2 µl template DNA. The PCR program was 35 cycles of 94°C for 30 s, 50°C for 1 min, and 72°C for 2 min, followed by a final extension at 72°C for 10 min as in[24]. Reaction products were examined by agarose gel electrophoresis. PCR products were directly cleaned with the Wizard SV Gel and PCR Clean-up System (Promega, Madison, USA) and analyzed by Sanger sequencing using four different primers: 18S-F[22], LABY-A[23], LABY-Y[23] and LABY-ARev[24]. Chromatograms were cleaned and assembled using DNASTAR software, with the Sanger Analysis and Assembly program. Assembled sequences have been deposited on NCBI with accession numbers MZ562737, MZ562738, MZ562739, MZ562740, MZ562741.

For phylogeny, obtained sequences were blasted on NCBI. 50 similar sequences were obtained, and *Oblongichytrium* and *E. huxleyi* 18S sequences were chosen as outgroup. We generated an alignment in mafft, keeping only sequences longer than 1000 bp, leaving 32 sequences in the final alignment. A neighbor-joining tree was performed on conserved sites (866 bp) with Jukes-Cantor model and 1000 bootstraps. The tree was exported in Newick format, and edited in Illustrator.

***Paired dilution experiment***

Phytoplankton growth and microzooplankton grazing rates were estimated using the dilution method[25,26]. A slightly modified version of the method was used with only one low dilution level (20%) and an undiluted treatment used[27]. Rates calculated using this method are considered conservative but accurate when compared with those using multiple dilution levels and a linear regression. Water from bags 1-4 was collected using a peristaltic pump at ~1m depth and mixed into a 20 L clean carboy. Water was screened through a 200 µm mesh to remove larger mesozooplankton. The collected water was shaded with black plastic and returned to shore. Dilution experiments were set-up in a temperature-controlled room, set to ambient water temperature (±2°C). Particle-free diluent (FSW) was prepared by gravity filtering whole seawater (WSW) through a 0.45 µm inline filter (PALL Acropak™ Membrane capsule) into a clean carboy. To the FSW, WSW was gently siphoned at a proportion of 20%. The 20% dilution and 100% WSW treatments were prepared in single carboys and then siphoned into triplicate 1.2 L Nalgene™ incubation bottles. To control for nutrient limitation, additional triplicate bottles of 100% WSW were incubated without added nutrients (10 µM nitrate and 1 µM phosphate). The incubation bottles were incubated for 24 hours in an outdoor tank maintained at *in-situ* water temperatures by a flow-through system of ambient seawater. Bottles could float freely, and the seawater inflow caused gentle agitation throughout the 24-hour period. A screen was used to mimic light conditions experienced within the mesocosm bags.

To quantify viral mortality, we used the paired dilution method[28] which involves setting up an extra low dilution level (20%) containing water filtered through a tangential flow filter (TFF) of 100 kDå to remove viral particles. During this experiment, TFF water was produced 1-2 days prior to the dilution experiment, to ensure the chemical composition of the water was as similar as possible, and experiments could be set up in a timely manner.

At T0 hours and T24 hours from all dilution experiments, sub-samples were taken for the determination of chlorophyll-*a* and flow cytometry. For chlorophyll-*a*, 100 – 150mL of seawater was filtered under low vacuum pressure through a 47mm Whatman GF/F filters (effective pore size 0.7 µm), and then extracted in 7mL of 97% methanol at 4°C in the dark for 12 hours. All chlorophyll readings were conducted on a Turner TD700 fluorometer[29]. Methanol blanks were included, and all samples were corrected for phaeophytin using a drop of 10% hydrochloric acid and then reading the sample again[30].

Water samples (2 x 1mL) for flow cytometry were taken at T0 and T24 of dilution experiments for the determination of phytoplankton abundances. Water samples were taken in triplicate from T0, and from each bottle at T24. Samples were immediately fixed in 20 µL of glutaraldehyde (final concentration <1%), gently inverted and then stored at 4°C for up to 2 hours. Samples were then flash frozen in liquid nitrogen and kept at -80°C until analysis. Samples were thawed and run at a high flow rate (104 – 108 µL $min^{-1}$) on a FACSCalibur (Becton Dickinson, East Rutherford, USA) for 1 – 5 minutes, based on the number of events triggered per second. Phytoplankton groups were differentiated into 4 groups; picoeukaryotes, nanoeukaryotes, *Synechococcus*, and *E. huxleyi* as explained above.

The apparent growth rates (*k*) of the total phytoplankton community (chlorophyll-*a*) and individual phytoplankton groups was calculated using the equation:

$$k = 1/t \ ln(C_t - C_0)$$

Where t = incubation time in days, $C_t$ and $C_0$ are the final and initial concentrations of chlorophyll-*a* or cell counts respectively.

Grazing and growth rates were calculated as in equation 4 and 5 of Morison and Menden-Deuer (2017). Grazing (*g*) was calculated as:

$$g = (k_d - k_1) / (1 - x)$$

Where, $k_d$ is the average growth rate in the diluted treatment (20%) and $x$ is the fraction of WSW, and $k_1$ is the average growth rate in 100% WSW with nutrient addition. Once grazing rates were calculated, the intrinsic growth rate ($\mu$) is calculated using $k_1$, which is the average growth rate without nutrients added:

$$\mu = g + k_1$$

Paired t-tests were conducted to determine significant differences ($p < 0.1$) between 100% WSW with and without nutrient additions. If no difference was found, the growth rates were pooled for calculations, otherwise calculated as above. Significant grazing rates were also determined through paired t-tests ($p < 0.1$) between 100% WSW and diluted treatments (20% WSW). Viral lysis was calculated as above for grazing, and if detected we also checked for a significant difference ($p < 0.1$) between diluted treatments with FSW and TFF waters to determine if the technique was sensitive enough to determine differences. On dates when viral lysis was determined, the intrinsic growth rate was calculated using both grazing and viral lysis rates. Results are shown in **Supplementary Fig. 3**.

***Estimated flux of organic carbon derived from E. huxleyi***

The estimated amount of organic carbon derived from *E. huxleyi* was calculated as follows. The volume V of an *E. huxleyi* cell was calculated based on a sphere of radius R=2.5µm using the formula $V = \frac{4}{3}\pi R^3 \sim 65.4498$ µm$^3$. The carbon content for one *E. huxleyi* was calculated by using a volume to carbon conversion factor of 220 fg C/µm$^3$ as in [31] leading to an estimate of 14,398 fg C/cell or 14.398 pg C/cell. We then estimated a loss of 19,050 *E. huxleyi* cells/ml/day, which corresponds to the difference in average abundances between day 17 (57000 cells/ml) and day 19 (18900 cells/ml). This corresponds to a loss of 274,281 pg C/ml/day or 274.3 ng C/ml/day or 274 µC/L/day.

***Transparent exopolymer (TEP) and Coomassie stainable particles (CSP).*** *TEP* concentration was determined following the spectrophotometric method[32]. Duplicate samples (50-200 mL) were filtered onto 25 mm diameter 0.4 µm pore size polycarbonate filters (DHI, San Francisco, USA) using a constant low filtration pressure (~150 mmHg). Immediately, the filters were stained with an Alcian Blue solution (500 µL, 0.02 %, pH 2.5) for 5 s, and rinsed with MilliQ water. Duplicate blanks (empty filters) were stained with every batch of samples and all filters were stored frozen (−20 °C) in 2mL Eppendorf tubes until further processing. Dye extraction of all filters was done by soaking in 5 mL of 80 % sulfuric acid for 3 h, shaking them intermittently. Absorbance of samples and blanks was measured against MilliQ water at 787 nm using the Varian Cary 100 Bio, and the mean absorbance of daily blank filters was subtracted from each batch of samples. The staining solution was calibrated following the original method of[32] with a xanthan gum standard and TEP concentration is reported in micrograms of xanthan gum equivalents per liter (µg XG eq/L).

CSP concentration was determined by spectrophotometry following[33]. Duplicate samples (60-200 mL) were filtered onto 25 mm diameter 0.4 µm pore size polycarbonate filters (DHI) using a constant low filtration pressure (~150 mmHg). The samples were immediately stained with 1 mL of Coomassie Brilliant Blue (CBB-G 250) solution (0.04 %, pH 7.4) for 30 s, prepared daily with filtered 0.2 µm fjord water collected at the beginning of the experiment, and rinsed three times with MilliQ water. Duplicate blanks (empty filters) were stained with every batch of samples and all filters were stored frozen (−20 °C) in 2mL Eppendorf tubes until further processing. Dye extraction of all filters was performed by soaking them in 4mL of extraction solution (3% SDS in 50% isopropyl alcohol) for 2 h at 37 ºC, shaking them every 30 minutes. Absorbance of samples and blanks was measured against MilliQ water at 615nm (Varian Cary 100 Bio), and the mean absorbance of daily blank filters was subtracted from each batch of samples. The staining solution was calibrated with a bovine serum albumin standard and CSP concentrations are expressed accordingly in micrograms of bovine serum albumin equivalents per liter (µg BSA eq/L).

***Dissolved organic carbon (DOC)***. For DOC determination, 30 mL samples of filtered sea water (GF/F, Whatman, Maidstone, UK) were collected in acid-cleaned polycarbonate bottles, and stored in the dark at −20 °C until analysis. They were analyzed with a TOC-LCSV (Shimadzu, Kyoto, Japan), with MilliQ water as a blank, potassium hydrogen phthalate as the calibration standard, and deep Sargasso Sea water as the reference (Hansell Laboratory. University of Miami, RSMAS). Each sample was injected repeatedly 4 to 5 times, until at least 3 reads yielded a relative standard deviation lower than 3%.

***Particulate organic carbon (POC) and nitrogen (PON), and Particulate inorganic carbon (PIC)***. For POC analyses, seawater (150-1000 mL) was filtered through combusted (4 h, 450 ºC) GF/F glass fiber filters (Whatman, Maidstone, UK) and filters were frozen at -20 ºC until processed. Prior to analysis, the filters were thawed in an HCl-saturated atmosphere for 48 h to remove inorganic compounds and dried at 80ºC for 24h[34]. Then the filters were dried and analyzed with an elemental analyzer (Perkin-Elmer 2400 CHN, Perkin-Elmer, Waltham, USA). For total particulate carbon (TPC) and PON the same procedure was followed except for the filter exposure to HCl-saturated atmosphere. PIC concentration was obtained subtracting POC from TPC values.

***Chlorophyll a (Chl a).*** Samples (100-250 mL) for fluorometric Chl *a* analysis were filtered on glass fiber filters (GF/F, 25 mm diameter, Whatman, Maidstone, UK) and stored at -20 ºC until analysis. Pigments were extracted with 90% acetone at 4 ºC in the dark for 24 hours. Fluorescence of extracts was measured, and corrected for phaeopigments, with a calibrated Turner Designs fluorometer[35].

***Polysaccharide analysis of particulate organic matter***. A peristaltic pump and tubings with a 200 µm mesh were used to sample between 25 to 100 L of water from the enclosures, which was subsequently filtered through pre-combusted 0.7 µm GF/F filters (Whatman, Maistone, UK) to harvest particular organic matter (POM).
Polysaccharide extraction: For the POM samples, 7 circular filter sections (11.2 mm diameter) were punched out from each GF/F filter and transferred into a 2 ml tube. Polysaccharides were sequentially extracted with: MilliQ water, 50 mM EDTA pH 7.5 and 4 M NaOH with 0.1% w/v $NaBH_4$. For each of the extracting solvents the following was performed: 400 µl of solvent were added to the tubes containing the filter pieces, vortexed them briefly and tubes were then incubated 2 h at 650 rpm (MilliQ at 60 °C and the other two solvents at room temperature). Samples were spun down at 6000 x g for 10 min at 15 °C. Extracts (supernatants) were collected in 1.5 ml tubes. The pellets and filter pieces were resuspended in the next extracting solvent using the same extraction procedure as depicted above.
Carbohydrate microarray analysis: All POM polysaccharide extracts were added into wells of 384-microwell plates. For each extract a 2-fold dilution followed by a 5-fold dilution was performed in printing buffer (55.2% glycerol, 44% water, 0.8% Triton X-100). Plates containing the samples were spun down at 3500 x g for 10 min at 15 °C to get rid of bubbles. The content of the plates was printed onto nitrocellulose membrane with a pore size of 0.45 µm (Whatman, Maidstone, UK) using a microarray robot (Sprint, Arrayjet, Roslin, UK) under controlled conditions of 20 °C and 50% humidity. A printing replicate was included for each sample. Once printed, each single microarray was individually probed with one glycan-specific monoclonal antibody. Microarray probing was performed as described previously[36]. The

developed arrays were scanned at 2400 dots per inch and binding of each probe (probe signal intensity) against each spotted sample was quantified using the software Array-Pro Analyzer 6.3 (Media Cybernetics, Rockville, USA). Array data analysis was performed as described previously[36]. Briefly, for each extract the mean antibody signal intensity was calculated. The highest mean signal intensity detected in the data set was set to 100 and all other values were normalized accordingly. Controls for the extraction solvents indicated no unspecific binding to any of the probes and controls for the anti-rat alkaline phosphatase-conjugated secondary antibody presented no unspecific binding to any of the samples and a cut-off of 5 arbitrary units was applied.

**References**


1. Rousseeuw, P. J. Silhouettes: A graphical aid to the interpretation and validation of cluster analysis. *J. Comput. Appl. Math.* **20**, 53–65 (1987).
2. Ramond, P. *et al.* Coupling between taxonomic and functional diversity in protistan coastal communities. *Environ. Microbiol.* **21**, 730–749 (2019).
3. Grasshoff, K. *Methods of seawater analysis*. (Verlag chemie, 1983).
4. Vincent, F. *et al.* AQUACOSM VIMS-Ehux – Core data. *Dryad* (2020). doi:Https://doi.org/10.5061/dryad.q573n5tfr
5. Picheral, M., Colin, S. & Irisson, J.-O. EcoTaxa, a tool for the taxonomic classification of images. (2017). Available at: http://ecotaxa.obs-vlfr.fr.
6. Caporaso, J. G. *et al.* Global patterns of 16S rRNA diversity at a depth of millions of sequences per sample. *Proc. Natl. Acad. Sci. USA* **108**, 4516–4522 (2011).
7. Parada, A. E., Needham, D. M. & Fuhrman, J. A. Every base matters: Assessing small subunit rRNA primers for marine microbiomes with mock communities, time series and global field samples. *Environ. Microbiol.* **18**, 1403–1414 (2016).
8. Apprill, A., Mcnally, S., Parsons, R. & Weber, L. Minor revision to V4 region SSU rRNA 806R gene primer greatly increases detection of SAR11 bacterioplankton. *Aquat. Microb. Ecol.* **75**, 129–137 (2015).
9. Callahan, B. J. *et al.* DADA2: High-resolution sample inference from Illumina amplicon data. *Nat. Methods* **13**, 581–583 (2016).
10. Wang, Q., Garrity, G. M., Tiedje, J. M. & Cole, J. R. Naïve Bayesian classifier for rapid assignment of rRNA sequences into the new bacterial taxonomy. *Appl. Environ. Microbiol.* **73**, 5261–5267 (2007).
11. Stoeck, T. *et al.* Multiple marker parallel tag environmental DNA sequencing reveals a highly complex eukaryotic community in marine anoxic water. *Mol. Ecol.* **19**, 21–31 (2010).
12. Piredda, R. *et al.* Diversity and temporal patterns of planktonic protist assemblages at a Mediterranean Long Term Ecological Research site. *FEMS Microbiol. Ecol.* **93**, (2017).
13. Guillou, L. *et al.* The Protist Ribosomal Reference database (PR2): a catalog of unicellular eukaryote small sub-unit rRNA sequences with curated taxonomy. *Nucleic Acids Res.* **41**, D597-604 (2013).
14. McMurdie, P. J. & Holmes, S. phyloseq: An R Package for Reproducible Interactive Analysis and Graphics of Microbiome Census Data. *PLoS One* **8**, e61217 (2013).
15. Laber, C. P. *et al.* Coccolithovirus facilitation of carbon export in the North Atlantic. *Nat. Microbiol.* **3**, 537–547 (2018).
16. Bai, M. *et al.* Molecular Detection and Spatiotemporal Characterization of Labyrinthulomycete Protist Diversity in the Coastal Waters Along the Pearl River Delta. *Microb. Ecol.* **77**, 394–405 (2019).

17. Martin, K. *et al.* The biogeographic differentiation of algal microbiomes in the upper ocean from pole to pole. *Nat. Commun.* **12**, 1–15 (2021).
18. Seddiki, K. *et al.* Sequencing, de novo assembly, and annotation of the complete genome of a new thraustochytrid species, strain CCAP_4062/3. *Genome Announc.* **6**, (2018).
19. Liu, B. *et al.* Draft genome sequence of the docosahexaenoic acid producing thraustochytrid Aurantiochytrium sp. T66. *Genomics Data* **8**, 115 (2016).
20. Kimura, H., Fukuba, T. & Naganuma, T. Biomass of thraustochytrid protoctists in coastal water. *Mar. Ecol. Prog. Ser.* **189**, 27–33 (1999).
21. Christian, J. R. & Karl, D. M. Microbial community structure at the U.S.-Joint Global Ocean Flux Study Station ALOHA: Inverse methods for estimating biochemical indicator ratios. *J. Geophys. Res.* **99**, 269–283 (1994).
22. Medlin, L., Elwood, H. J., Stickel, S. & Sogin, M. L. The characterization of enzymatically amplified eukaryotic 16S-like rRNA-coding regions. *Gene* **71**, 491–499 (1988).
23. Stokes, N., Ragone Calvo, L., Reece, K. & Burreson, E. Molecular diagnostics, field validation, and phylogenetic analysis of Quahog Parasite Unknown (QPX), a pathogen of the hard clam Mercenaria mercenaria. *Dis. Aquat. Organ.* **52**, 233–247 (2002).
24. Collado-Mercado, E., Radway, J., Ecology, J. C.-A. M. & 2010, U. Novel uncultivated labyrinthulomycetes revealed by 18S rDNA sequences from seawater and sediment samples. *Aquat. Microb. Ecol.* **58**, 215–228 (2010).
25. Landry, M. R. & Hassett, R. P. Estimating the grazing impact of marine micro-zooplankton. *Mar. Biol. 1982 673* **67**, 283–288 (1982).
26. Landry, M. R., Kirshtein, J. & Constantinou, J. A refined dilution technique for measuring the community grazing impact of microzooplankton, with experimental tests in the central equatorial Pacific. *Mar. Ecol. Prog. Ser. Ecol. Prog. Ser.* **120**, 53–63 (1995).
27. Morison, F. & Menden-Deuer, S. Doing more with less? Balancing sampling resolution and effort in measurements of protistan growth and grazing-rates. *Limnol. Oceanogr. Methods* **15**, 794–809 (2017).
28. Evans, C., Archer, S. D., Jacquet, S. & Wilson, W. H. Direct estimates of the contribution of viral lysis and microzooplankton grazing to the decline of a Micromonas spp. population. *Aquat. Microb. Ecol.* **30**, 207–219 (2003).
29. Jespersen, A. & Christoffersen, K. Measurements of chlorophyll—a from phytoplankton using ethanol as extraction solvent. *Arch. Fur Hydrobiol.* (1987).
30. Holm-Hansen, O. & Riemann, B. Chlorophyll a Determination: Improvements in Methodology. *Oikos* **30**, 438 (1978).
31. Simó, R. *et al.* Annual DMSP contribution to S and C fluxes through phytoplankton and bacterioplankton in a NW Mediterranean coastal site. *Aquat. Microb. Ecol.* **57**, 43–55 (2009).
32. Passow, U. & Alldredge, A. L. A dye-binding assay for the spectrophotometric measurement of transparent exopolymer particles (TEP). *Limnology and Oceanography* **40**, 1326–1335 (1995).
33. Cisternas-Novoa, C., Lee, C. & Engel, A. A semi-quantitative spectrophotometric, dye-binding assay for determination of Coomassie Blue stainable particles. *Limnol. Oceanogr. Methods* **12**, 604–616 (2014).
34. Hansen, P., Koroleff, F., Grasshoff, K., Kremling, K. & Ehrhardt, M. *Determination of nutrients, Methods of Seawater Analyses*. (Wiley-Vch, 1999).
35. Holm-Hansen, O., Lorenzen, C. J., Holmes, R. W. & Strickland, J. D. H. Fluorometric Determination of Chlorophyll. *ICES J. Mar. Sci.* **30**, 3–15 (1965).

36. Vidal-Melgosa, S. *et al.* A new versatile microarray-based method for high throughput screening of carbohydrate-active enzymes. *J. Biol. Chem.* **290**, 9020–9036 (2015).
